# Neotelomeres and Telomere-Spanning Chromosomal Arm Fusions in Cancer Genomes Revealed by Long-Read Sequencing

**DOI:** 10.1101/2023.11.30.569101

**Authors:** Kar-Tong Tan, Michael K. Slevin, Mitchell L. Leibowitz, Max Garrity-Janger, Heng Li, Matthew Meyerson

**Affiliations:** Department of Medical Oncology, Dana-Farber Cancer Institute, Boston, MA 02215, USA; Cancer Program, Broad Institute of MIT and Harvard, Cambridge, MA 02142, USA; Department of Genetics, Harvard Medical School, Boston, MA 02215, USA; Center for Cancer Genomics, Dana-Farber Cancer Institute, Boston, MA 02215, USA; Department of Data Sciences, Dana-Farber Cancer Institute, Boston, MA 02215, USA; Department of Biomedical Informatics, Harvard Medical School, Boston, MA 02215, USA

**Keywords:** Telomere, long-read sequencing, neotelomeres, arm fusions, repetitive elements

## Abstract

Alterations in the structure and location of telomeres are key events in cancer genome evolution. However, previous genomic approaches, unable to span long telomeric repeat arrays, could not characterize the nature of these alterations. Here, we applied both long-read and short-read genome sequencing to assess telomere repeat-containing structures in cancers and cancer cell lines. Using long-read genome sequences that span telomeric repeat arrays, we defined four types of telomere repeat variations in cancer cells: neotelomeres where telomere addition heals chromosome breaks, chromosomal arm fusions spanning telomere repeats, fusions of neotelomeres, and peri-centromeric fusions with adjoined telomere and centromere repeats. Analysis of lung adenocarcinoma genome sequences identified somatic neotelomere and telomere-spanning fusion alterations. These results provide a framework for systematic study of telomeric repeat arrays in cancer genomes, that could serve as a model for understanding the somatic evolution of other repetitive genomic elements.

## Introduction

Cancer is driven by alterations to the genome. The continued invention and application of new methods has enabled the characterization of genomic alterations in cancer with much greater scale and resolution. The development of massively parallel short-read sequencing over the past fifteen years has greatly accelerated our efforts to characterize the cancer genome by enabling the detailed and rapid characterization of somatic and germline variants in tens of thousands of samples^1–7^, and led directly to the discovery of many cancer driving genetic alterations that are now being targeted by emerging therapeutics. The recent development and application of linked-read genome sequencing of long molecules with barcoded short-reads then facilitated the characterization of more complex structural variations, and of genomic alterations at the haplotype-level in cancer^8–12^.

Despite these advances in genome technology, the identification and characterization of somatic alterations at repetitive elements, which constitute approximately half the human genome^13–15^, still remain significant challenges. Repetitive elements and duplicated sequences in the human genome are typically 100 to 8000 bp in size^15^, although centromeres are much longer arrays of repetitive elements, and can be broadly classified into three main classes. First, repetitive elements include tandem repeats of specific DNA sequences^15,16^, including short tandem repeats (1-6 bp repeat unit) in the form of microsatellites, and longer repeat units forming minisatellites^16^. Telomeres and centromeres, which are key structures in a chromosome, are largely comprised of long tandem repeats^15^. Second, repetitive elements include interspersed repeats, identical or nearly identical sequences spread out across the human genome^15^, including short interspersed nuclear elements (SINEs; typically 100-300 bp in length) such as Alu repeats, and long interspersed nuclear elements (LINEs, typically >300 bp in length) such as L1 repeats^15^. Third, “low copy repeats”, or segmental duplicates, also occur in the genome. These large repetitive sequences are blocks of DNA that are 1-400 kilobases in size, occur as at least two copies, share high sequence similarity (>90%)^17,18^, and are potential hotspots of chromosomal rearrangements and instability^19,20^. Although sophisticated computational methods have been developed to infer somatic alterations in repetitive regions using short-reads, comprehensive characterization of somatic alterations in these regions still cannot be completely achieved.

Telomeres are a salient example of highly repetitive structures of particular importance in cancer that cannot yet be readily resolved by current sequencing methodologies. Human telomeres, which act as protective caps on the ends of chromosomes are composed of ∼2-10 kb (TTAGGG)_n_ tandem repeats^21,22^. Somatic integration of telomeric sequences into non-telomeric DNA in tumor samples has also been observed^23^, though the origin and structures of these sequences remain unclear. As the short-read sequencing that is typically performed, such as 2 x 150 bp paired reads, is unable to fully span the 2 kb – 10 kb long highly repetitive telomeres, much remains unknown about telomere structures in cancer.

The study of telomere structure is important in cancer genomics because telomere maintenance is crucial in cancer pathogenesis. Cancer cell immortality requires a mechanism to activate telomerase or otherwise maintain telomeres, and is a key “hallmark of cancer”^24,25^. Telomerase, the enzyme which adds telomeric repeats to the ends of chromosomes, has been estimated to undergo reactivation in as many as 90% of human cancers and was shown experimentally to be critical for malignant transformation^26–31^. The reactivation of telomerase activity in cancer is driven in part by promoter mutations, amplifications and translocations in the telomerase catalytic subunit gene, *TERT*^32–35^, and also by amplification of the RNA component of telomerase, *TERC*, in cancer^35,36^. In some cancer types, genetic inactivation of the *ATRX* and *DAXX* genes are also associated with telomere elongation, independent of telomerase, by the alternative lengthening of telomeres (ALT) pathway^37,38^.

The emergence of long-read genome sequencing now makes it possible to analyze somatic alterations in highly repetitive regions, such as telomeric repeats, with greater precision and detail. Recently, the first telomere-to-telomere human genome was assembled using long-reads that can span large, complex, or repetitive genomic sequences, including telomeric repeats. This assembly relied upon PacBio high-fidelity (HiFi) sequencing, which can generate long reads with an accuracy of 99.8% and an average length of 13.5 kb^39^, as well as ultra-long-read nanopore sequencing, with can generate reads of over 100 kb^40^, Using long reads, the repetitive telomeres can be spanned and mapped uniquely to the human genome. However, long-read sequencing is still significantly more expensive than short-read sequencing. Given that high-coverage short-read genome sequences are now widely available, a cost-effective strategy at this time is to leverage short-read sequencing datasets to identify samples with potentially interesting telomeric alterations *in silico*, and to subject these samples to more detailed analysis by long-read sequencing.

Here, we explored the structure of previously unresolved telomeric events in the cancer genome. We used large databanks of short-read genome sequencing datasets to identify candidate telomeric alterations in the genome of 326 cancer cell line and 95 primary lung adenocarcinoma samples using TelFuse, a computational method to profile ectopic intra-chromosomal telomeric repeat sites. Then, using PacBio HiFi and Nanopore long-read genome sequencing in three cell lines with high numbers of putative telomeric variants, we resolved the structure of these alterations in combination with spectral karyotyping, copy number and allelic ratio analysis. Long-read genome sequencing of these samples led directly to the discoveries of neotelomeres, telomere-spanning chromosomal arm fusion events, and complex telomeric alterations that were not previously resolvable using short-read genome sequencing. These findings also validate recent experimental observations on neotelomere formation^41^. Our study creates a framework that can be applied to the examination of other highly repetitive sequences that are likely to be of biological significance in disease, including centromere arrays, transposable element insertions, and microsatellite repeats.

## Results

### Identification of ectopic telomeric repeat sequences

Telomeric repeat arrays within cancer genomes can be found at their original position at chromosomal termini (**Figure 1A**), or at new positions within the genome (**Figure 1B**). Telomeric repeats at new genomic locations may be in the same orientation as the original telomeric repeat, with reference to the adjacent chromosomal sequence (i.e. standard orientation), or in an inverted orientation (**Figure 1B**). Significantly, telomeric repeats oriented in different directions may represent different chromosome structures and may originate via highly distinct biological processes.

**Figure 1.**
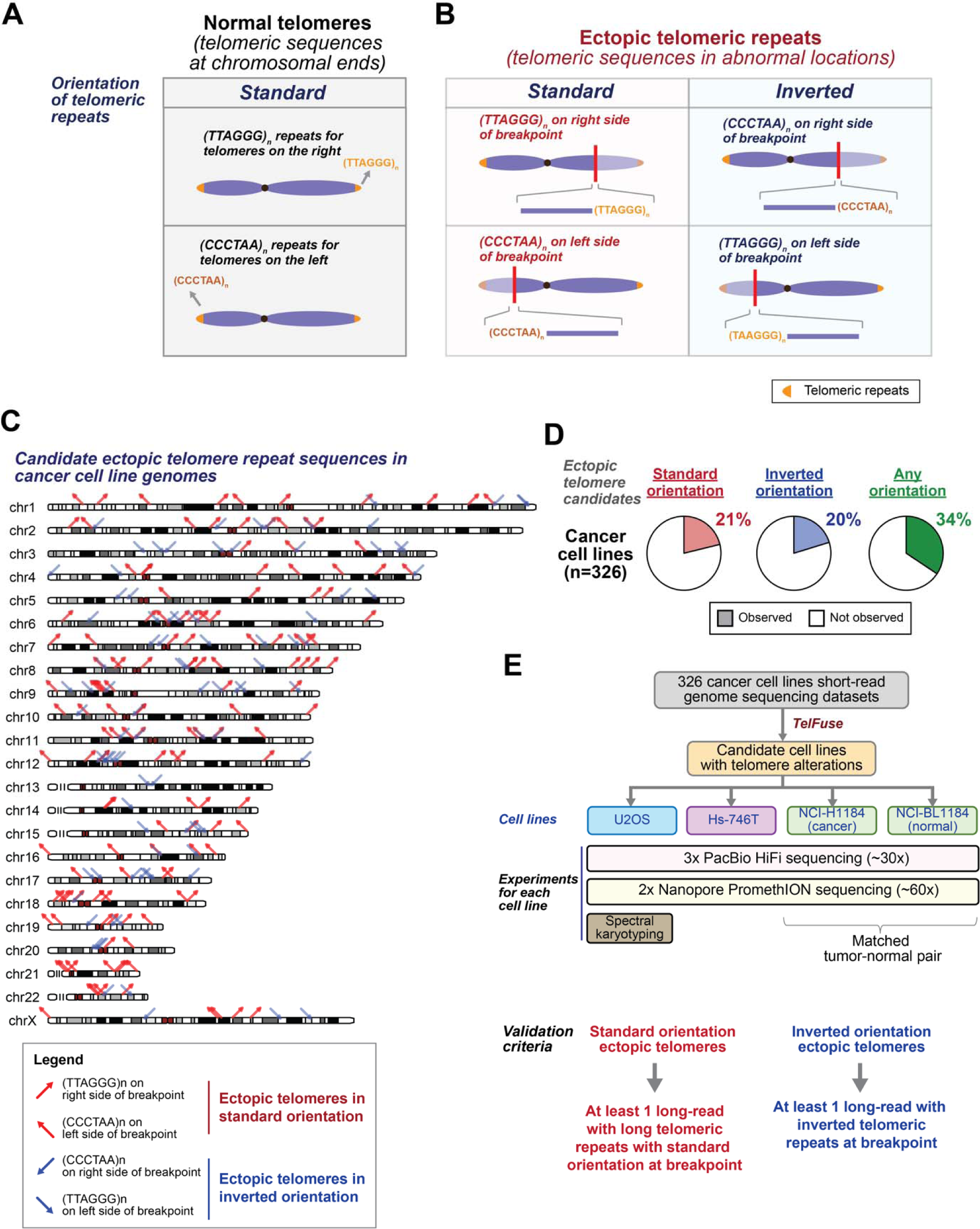
Classes of ectopic telomeric repeats found in cancer cell genomes. **(A)** Schematic of sequence and positions of normal telomeres at chromosomal termini. **(B)** Schematic of ectopic telomeric repeats found at abnormal locations away from chromosomal termini. Standard orientation: (TTAGGG)_n_ on the right side of a breakpoint and (CCCTAA)_n_ on the left side of the breakpoint in the 5’ to 3’ direction (same as normal telomere in Fig. 1A). Inverted orientation: (CCCTAA)_n_ on the right side of a breakpoint and (TTAGGG)_n_ on the left side of the breakpoint in the 5’ to 3’ direction. Note that faded chromosomal segment is not part of derivative chromosome. **(C)** Genome-wide localization of ectopic telomeric repeats in cancer cell line genomes (n=326) identified using short-read genome sequencing. Red: ectopic telomeric sequences in the standard orientation. Blue: ectopic telomeric sequences in the inverted orientation. Position of telomeric repeats relative to the breakpoint is indicated by arrows oriented in different directions. **(D)** Percentage of cancer cell lines in the CCLE with ectopic telomeric sequences in either orientation. Total sample number as indicated. **(E)** Flow-chart of long-read genome sequencing and cytogenetic analyses in cancer cell lines, with the indicated validation criteria.

We developed the analytic method, TelFuse, to identify ectopic telomeric repeats within the cancer genome, and to estimate telomere length of each chromosomal arm with long-read sequencing respectively **(Figure S1A)**. TelFuse identifies ectopic telomeric repeat sequences (TTAGGG)_n_ or (CCCTAA)_n_ that are absent from the germline and mapped to intrachromosomal regions (i.e. at least 500 kb from chromosomal ends) (**Methods, Figure S1A-B)**. TelFuse begins by identifying read pairs that contain at least 2 perfect consecutive telomeric repeats (at least 12 base pairs of telomere sequence) with adjacent sequences that map to intra-chromosomal sites. Paired read sequences that are fully aligned to the reference genome are removed, eliminating telomeric repeats in the reference, which include ancient chromosome fusion events^42,43^. To ensure the specificity of our calls, we also developed a series of filters **(Figure S1A-B, Methods)** to remove spurious sites caused by artefacts induced during the mapping process (**Figure S1A-B**), assessed by a variety of quality control metrics (**Methods**). Those sites that pass all filters and are at least 500 kb from the GRCh38 reference genome chromosome terminus, a sufficient distance to avoid sub-telomere sequences^44–46^, are considered candidate sites of ectopic telomere sequence.

### Frequency and genome-wide distribution of candidate ectopic telomere repeat sequences in cancer cell lines inferred from short-read genome sequencing

To assess the landscape of ectopic telomere repeats in cancer, we began by analysis of cancer cell line data, which allows assessment of multiple cancer types and which provides high sequencing depth due to 100% cancer cell purity. We applied TelFuse to whole genome sequencing datasets from 326 cancer cell line DNA specimens from the Cancer Cell Line Encyclopedia (CCLE)^7^, and detected 240 candidate ectopic intra-chromosomal telomeric repeat sequence sites in 34% of cell lines (112/326) (**Figure 1C-D and Table S1 and S2**). Analysis of the orientation of the telomere repeats further defined these candidates as corresponding to 149 candidate sites with telomeric repeat sequences in the standard orientation, and 91 candidate sites with telomeric repeat sequences in the reverse orientation (**Figure 1C-D**). An additional 42 candidate sites with telomeric repeat sequences within softclipped sequences, but not on the first 12 base-pairs, were also detected **(Table S3)**; these were not analyzed in depth. These data indicate that genomic events involving telomeric repeat sequences can be readily detected in cancer cell lines from short-read genome sequencing using TelFuse.

### Validation of putative ectopic telomeres by long-read sequencing

Although short-read sequencing (2 x 101bp for the CCLE dataset^7^) can detect ectopic telomeric sequences, the length and repetitive nature of these sequences, which can span 10 kb in length^21,22^, renders their structures indecipherable based on short-read data alone, and cannot distinguish between possible modes of generation of these sites. Therefore, we decided to perform high-depth long-read sequencing of selected cell line genomes.

We selected the U2-OS osteosarcoma cancer cell line, with 55 candidate telomeric repeat sites from short read sequencing (46 in the standard orientation and 9 in the inverse orientation), the Hs-746T gastric carcinoma cell line with 6 candidate events (1 standard and 5 inverse orientation), and the NCI-H1184 small cell lung cancer cell line with 6 candidate events (5 standard and 1 inverted orientation) **(Figure S1C-E)**, together with its matched normal sample (NCI-BL1184). These samples were selected due to the high frequencies of ectopic telomeric events **(Figure S1C-E)**. Notably, the U2-OS cell line was found to be highly rearranged, with ectopic telomeric sites found near regions with changes in sequencing coverage and allelic ratios **(Figure S2)**. We then performed PacBio HiFi and Oxford Nanopore long-read genome sequencing (**Figure 1E**). We achieved a median genomic coverage of 49x, 62x, 65x and 73x for the U2OS, Hs-746T, NCI-BL1184 and NCI-H1184 cell lines respectively with Nanopore long-read genome sequencing **(Figure S3, Table S4 and S5).** With PacBio HiFi sequencing, we achieved a median genomic coverage of 19x, 20x, 19x, and 23x for the same four cell lines using high quality PacBio HiFi reads, and a median coverage of 29x, 31x, 33x and 36x when all PacBio reads were considered **(Figure S3, Table S4 and S5)**. Nanopore sequencing data had a median read length of 6-13 kb (N50: 18-21 kb), while the PacBio HiFi data had a median read length of 15-17 kb (N50: 16-19 kb) **(Figure S3, Table S4 and S5)**. In parallel, to assess chromosomal scale structures of these events, we also performed spectral karyotyping.

Long-read sequencing of cancer cell line genomes revealed two major types of structural alterations containing telomere repeat sequences that comprised either telomeric repeat sequences of > 1 kb flanked on one end by chromosomal sequence with no other flanking DNA (seen in 46 of 51 examples sequenced) or telomeric repeat sequences of at least few hundred base-pairs flanked on both sides by chromosomal sequence (seen in 12 of 15 candidate events sequenced). Telomeres flanked on only one-end with chromosomal sequence are consistent with neotelomere structures, which might be generated through telomerase activity^41^. Telomeres flanked on both sides with chromosomal sequence are likely to be sites of chromosome fusion or other translocation events.

### Neotelomeres in cancer revealed by long-read genome sequencing

Long-read sequencing analyses demonstrated that the ectopic telomere repeat sequences in the standard orientation were long and unbounded and therefore consistent with neotelomere addition. For example, a candidate ectopic telomere repeat sequence site adjacent to sequence from chrX:103,320,553 in the U2-OS osteosarcoma cell line was observed to contain at least seven tandem (TTAGGG)_n_ repeats in the short-read sequencing data (**Figure 2A**), and a reduction in sequencing coverage, corresponding to the position of the telomeric repeats, at this chromosomal position (**Figure 2B**). Upon analysis of this region in long-read sequencing data sets, both PacBio HiFi and Oxford Nanopore, long telomeric repeats of ∼3-10kb in the standard orientation could be readily observed (**Figure 2C**), where the variation in telomere length between reads might be explained by active telomere sequence loss or telomere maintenance after DNA replication, in different cells across the population. These data support a model where breakage of the chrXq arm was capped by generation of novel telomeric sequence representing a neotelomere (**Figure 2D**).

**Figure 2.**
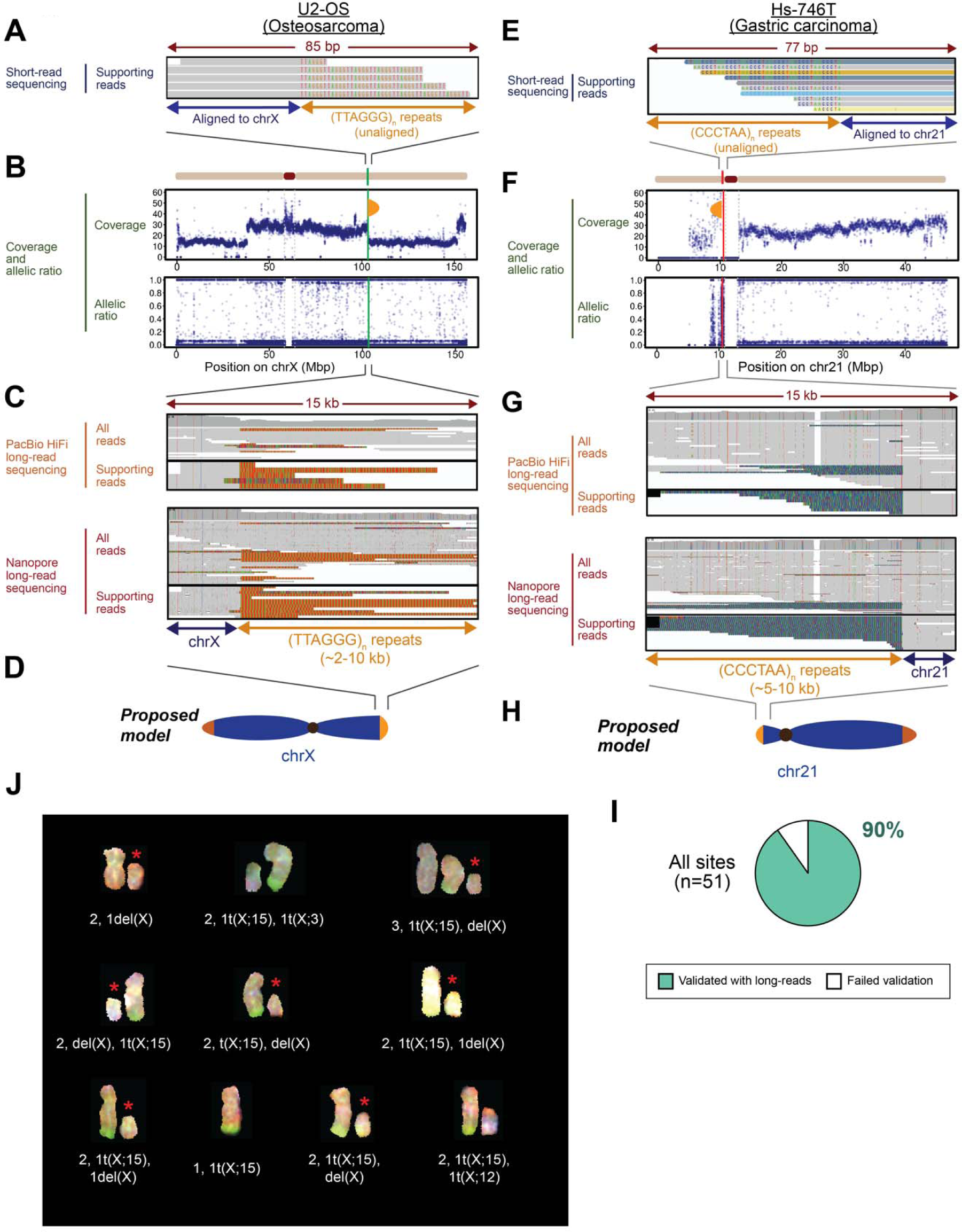
Neotelomeres in cancer genomes revealed by long-read genome sequencing. **(A-H)** Genomic analysis of telomere repeat alterations in the standard orientation that were detected **(A-D)** in the U2-OS osteosarcoma cell line at chrX:103,320,553, and **(E-H)** in the Hs-746T cell line at chr21:10,547,397. **(A)** IGV screenshots of short-read genome sequencing data. Ectopic telomeric repeats (TTAGGG)_n_ are shown in color. **(B)** Sequencing coverage and allelic ratios of chromosome X. Orange semi-oval: site of the neotelomeric event. **(C)** IGV screenshots depicting long telomeric repeat sequences (TTAGGG)_n_ with PacBio HiFi and Nanopore long-read sequencing at the site shown in (**A**). **(D)** Schematic of neotelomere location on chromosome Xq. **(E)** IGV screenshots of short-read genome sequencing data. Ectopic telomeric repeats (CCCTAA)_n_ are shown in color. **(F)** Sequencing coverage and allelic ratios of chromosome 21. Orange semi-oval: site of the neotelomeric event. **(G)** IGV screenshots depicting long telomeric repeat sequences (CCCTAA)_n_ with PacBio HiFi and Nanopore long-read sequencing at the site shown in (**E**). **(H)** Schematic of neotelomere location on chromosome 21p. **(I)** Percentage of ectopic telomeric repeat sites in the standard orientation, found by short-read genome sequencing using TelFuse, that were validated by long-read genome sequencing. **(J)** Spectral karyogram of chrX in ten U2-OS single cells assessed by spectral karyotyping with corresponding karyotype labels. First label: total # of X chromosomes and their derivatives observed in given cell. Second label: karyotypes of the aberrant X chromosomes or derivatives. Asterisk (*): truncated X chromosome. See also Figure S4.

Another example of a neotelomere is seen in the Hs-746T cell line, within chromosome arm 21p at chr21:10,547,397 where an ectopic telomeric repeat site was observed. Short-read sequencing showed at least six tandem (CCCTAA)_n_ repeats (**Figure 2E**). At this location, fluctuation in both sequencing coverage and allelic ratios could be observed (**Figure 2F**). Analysis of both PacBio HiFi and Nanopore long-read genome sequencing data again revealed long telomeric repeats (∼5-10 kb) in the standard orientation with reference to the break point at this site (**Figure 2G**), lending support to the existence of a neotelomere which had likely formed following breakage of the chr21p arm (**Figure 2H**). Similar observations were made at other ectopic neotelomeric sites, such as chr7:24,302,169 in the U2-OS cell line **(Figure S4A-D)**, and chr1:214,460,753 in the NCI-H1184 small cell lung carcinoma cell line **(Figure S4E-H)**, further supporting the idea that these ectopic telomeric sites in the standard orientation detected by short-reads represent neotelomere addition events.

In all, among 51 sites predicted by TelFuse as containing standard orientation telomere repeat sequences in these three cancer cell lines using short-read genome sequencing data, 46 of these sites could be readily demonstrated to represent long telomere repeats suggestive of neotelomeres, using the long-read genome sequencing data (**Figure 2I, Table S6)**. No telomeric long reads could be found at the other 5 sites. Together, our results indicate that short telomeric repeats in the standard orientation, observed with short-read sequencing data, represent neotelomeres with long telomeric repeats as confirmed by long-read genome sequencing.

To assess the relationship between neotelomeres and chromosomal alterations, and to support our neotelomere calls, we performed spectral karyotyping of the U2-OS cancer cell line, with detailed karyotyping for ten randomly selected cells (representative cell shown in **Figure S5A)**. Integrative analysis of sequencing coverage, allelic ratios and long-read data inferred two copies of chromosome X in U2-OS cells, one complete copy and one truncated chromosome X. Concordant with a neotelomere detected by long-read genome sequencing data (**Figure 2A-D**), a shorter chromosome X with q-arm deletion was observed by spectral karyotyping in 7/10 cells assessed (**Figure 2J**), together with a full-length chromosome X in 10/10 cells karyotyped. Thus, the spectral karyotyping analysis confirms that neotelomeres identified by long-read sequencing can be correlated with chromosomal truncations observed by cytogenetics.

We also observed a significant level of chromosomal heterogeneity **(Figure S5B-C, Table S7)**. Heterogeneities we observed included slight variations in chromosome number between each cell (N=76-80) **(Table S7)** and heterogeneity in translocation events between cells that were concordant with a prior study^27^. Specifically, while a t(4;22) translocation could be observed in 10/10 cells assessed **(Figure S5C)**, a t(15;19) translocation was only observed in 6/10 cells assessed. This cellular heterogeneity might explain why long-read sequencing was unable to validate 5 of the 51 candidate sites that were detected in the population of cells sequenced by CCLE. Therefore, heterogeneity in tumor cell populations remains a complication in identifying ectopic telomeric events.

### Telomere repeat-spanning chromosomal arm fusions in cancer resolved by long-read genome sequencing

We next explored sites with ectopic telomeric repeat sequences found in the inverted orientation with respect to the breakpoint; long-read sequencing revealed that these sites largely represent chromosomal arm fusion events. At one candidate site at position chr4:30,909,846, we observed eight inverted telomeric repeats (CCCTAA)_n_ (∼48 bp) using short-read sequencing data (**Figure 3A**). At this position, a significant change in sequencing coverage and change in allelic ratio were also observed in support of the fusion event (**Figure 3B**). Analyzing this region with both PacBio HiFi and Nanopore long-read genome sequencing, we observed ∼650bp of inverted (CCCTAA)_n_ repeats after the breakpoint (**Figure 3C**), followed by 5-8 kb of sequences on chr22q sub-telomeres (**Figure 3C**). Individual long-reads that cover the whole event suggest that the inverted (CCCTAA)_n_ repeat sequences formed via the fusion of the chr22q arm with its short telomere to an intra-chromosomal site (**Figure 3D**).

**Figure 3.**
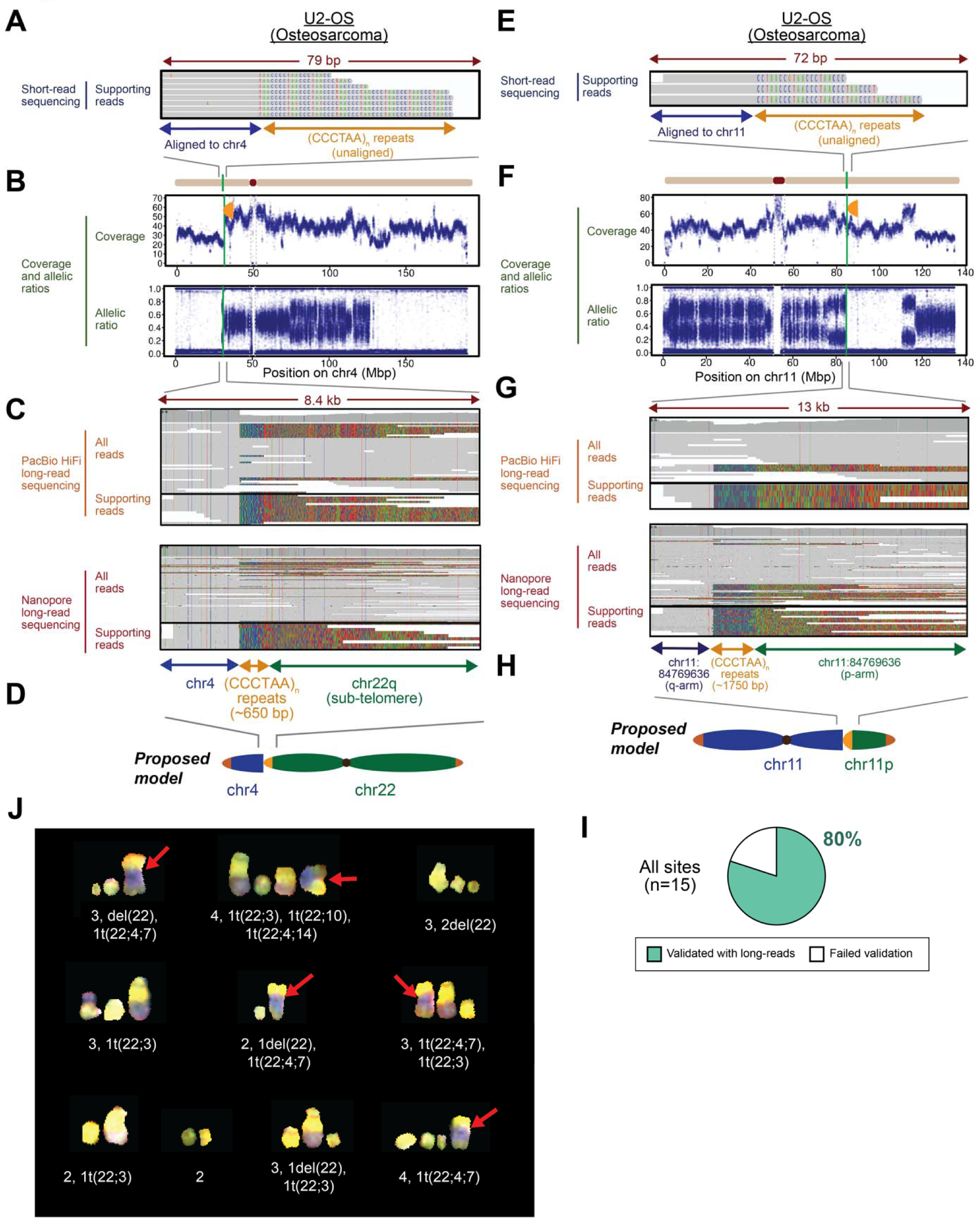
Chromosomal arm fusions in cancer genomes revealed by long-read genome sequencing. **(A-H)** Genomic analysis of telomere repeat alterations in the inverted orientation that were detected in the U2-OS osteosarcoma cell line **(A-D)** at the site chr4:30,909,846, and **(E-H)** at the site chr11:84,769,636. **(A)** IGV screenshots of short-read genome sequencing data. Ectopic telomeric repeats (CCCTAA)_n_ are shown in color. **(B)** Sequencing coverage and allelic ratios of chromosome 4. Orange semi-oval: site of the ectopic telomere repeat sequence. **(C)** IGV screenshots of PacBio HiFi and Nanopore long-read sequencing data at the site shown in (**A**). Ectopic telomeric repeats in the inverted orientation contained ∼650 bp of (CCCTAA)_n_ telomeric repeat sequences followed by chr22q sub-telomeric sequences, indicative of a chromosomal arm fusion event of chr22q to the site at chr4:30,909,846. **(D)** Schematic of telomere-spanning fusion event between chromosomes 22q-ter and 4p. **(E)** IGV screenshots of short-read genome sequencing data. Ectopic telomeric repeats (CCCTAA)_n_ are shown in color. **(F)** Sequencing coverage and allelic ratios of chromosome 11. Orange semi-oval: site of the ectopic telomere repeat sequence. **(G)** IGV screenshots of PacBio HiFi and Nanopore long-read sequencing at the site shown in (**E**). ∼1750 bp of (CCCTAA)_n_ telomeric repeat sequences are found sequences corresponding to chr11p (chr11:43,002,345), suggestive of a complex event consistent with the formation of a neotelomere on chr11p, followed by a chromosomal arm fusion event of this neotelomere to the site on chr11q (chr11:84,769,636). **(H)** Schematic telomere-spanning fusion event between chromosome arms 11q (with a predicted neotelomere) and 11p. **(I)** Percentage of new telomeric sites in the inverted orientation that were predicted by TelFuse from short-read genome sequencing, and then validated by long-read genome sequencing as telomere-spanning chromosome arm fusion events. **(J)** Spectral karyogram of chromosome 22 for which a chromosomal arm fusion was detected with chromosome 4. Ten U2-OS single cells assessed are as indicated. The fusion event between chromosome 22 (yellow) and chromosome 4 (blue) is indicated by a red arrow. See also Figure S6.

We also observed more complex fusion events, including evidence for the formation of a neotelomere followed by a subsequent chromosomal fusion. At chr11:84,769,636, five inverted ectopic telomeric repeats (CCCTAA)_n_ (∼30 bp) were detected at the breakpoint with short-read sequencing (**Figure 3E**). At this site, a drastic change in allelic ratios was observed despite minimal changes in copy number estimated from sequencing coverage (**Figure 3F**), suggesting changes to one of the parental chromosomes despite no overall changes in chromosomal number. Using both PacBio HiFi and Nanopore long-read sequencing data, we observed ∼1750 bp of inverted (CCCTAA)_n_ telomeric repeats at this site (**Figure 3G**). Surprisingly, we could further observe >5kb of sequences corresponding to an intra-chromosomal site on the chr11p arm, suggesting that the neotelomere was the consequence of multiple steps. It may have first formed on the centromeric side of the chr11p breakpoint (chr11p:43,002,345), which then subsequently fused to the breakpoint on chr11q at position 84,769,636 (**Figure 3H**).

To assess if telomere-spanning chromosomal fusions could be detected in other samples, we again examined long-read genome sequencing data of the Hs-746T gastric adenocarcinoma and NCI-H1184 lung adenocarcinoma cell lines. Inverted ectopic telomeric repeats that were identified using TelFuse were confirmed as sites of chromosomal arm fusion events with long-read data in the Hs-746T sample **(Figure S6)** at the sites chr11:79,325,679 and chr1:244,201,717, but not for the single candidate site in the NCI-H1184 sample **(Table S6)**. Again, the discrepancy between long- and short-read data in our study could be caused by heterogeneity in the cancer cell lines. Overall, across 15 inverted telomeric repeat sites predicted by TelFuse in these cell lines, 12 of these events (80%) could be validated as chromosomal arm fusion events using long-read genome sequencing (**Figure 3I, Table S6)**.

We further investigated chromosomal arm fusion events for their concordance with spectral karyotyping results of the U2-OS cells. Consistent with the t(4;22) fusion seen in long-read sequencing (**Figure 3A-D**), a fusion between chromosome 22 and chromosome 4 was observed by spectral karyotyping in 5/10 cells assessed (**Figure 3J**). As such, these results suggest that telomere-spanning chromosomal arm fusion events detected by long-read sequencing are concordant with the chromosomal scale observations.

### Length distribution of neotelomeres matches that of normal telomeres

Because short telomeres lead to chromosomal fusion events, we hypothesized that neotelomeres would have similar lengths to unaltered telomeres at chromosome ends, whereas fusion events, which might have resulted from telomere attrition, would be shorter. To assess telomere length, we developed an approach (TelSize) to estimate the length of telomeric repeats in long read sequences **(Methods)** that accounts for noise in telomeric long-reads which are interspersed with errors and/or *bona fide* deviations from the standard “TTAGGG” repeat motif **(Figure S7A)**.

Using TelSize, we can estimate the length of telomere repeat regions on single chromosomes. We applied the TelSize approach to establish the length of telomeres found at each of the chromosomal arms, and at intra-chromosomal telomeric sites. As the sub-telomeric region of the GRCh38 reference genome has not been fully assembled, we first assessed the reliability of assigning telomeric long reads to their respective arms for the CHM13 cell line for which the genome has been fully assembled (**Figure S7B**). TelSize was used to generate telomere length estimates for all of the cell lines with long read sequencing data **(Figure S8)**.

We then assessed the length of telomeres at each neotelomere, at each natural telomere found on each chromosomal arm, and each chromosomal arm fusion event. For example, in a site of neotelomere addition at position chrX:103,320,553 in DNA from U2-OS cells that was described in an earlier (**Figure 2A-D**), TelSize predicts a telomere length of at least 4988 bp from a single nanopore read (**Figure 4A**). In a site of chromosome arm fusion between positions chr4:30,909,846 and the chr22 telomeric end (**Figure 3A-D**) in DNA from U2-OS cells, TelSize predicts a telomere length of 632 bp from a single nanopore read (**Figure 4B**), with intra-chromosomal and sub-telomeric sequences flanking these sites. Most neotelomeres identified were multi-kilobasepair long with an average telomere length of ∼5kb in both the U2-OS and the Hs-746T cancer cell lines (**Figure 4C-D, Figure S9A-B)**. In contrast to neotelomeres and normal chromosomal arms, and consistent with our hypothesis, we see that telomeres at chromosomal arm fusion events tend to be relatively short and were largely only a few hundred base pairs long in U2-OS but longer in the small number of examples in Hs- 746T (**Figure 4E-F, Figure S9C-D)**, suggesting that chromosomal arms with short telomeres are more likely to undergo fusion events.

**Figure 4.**
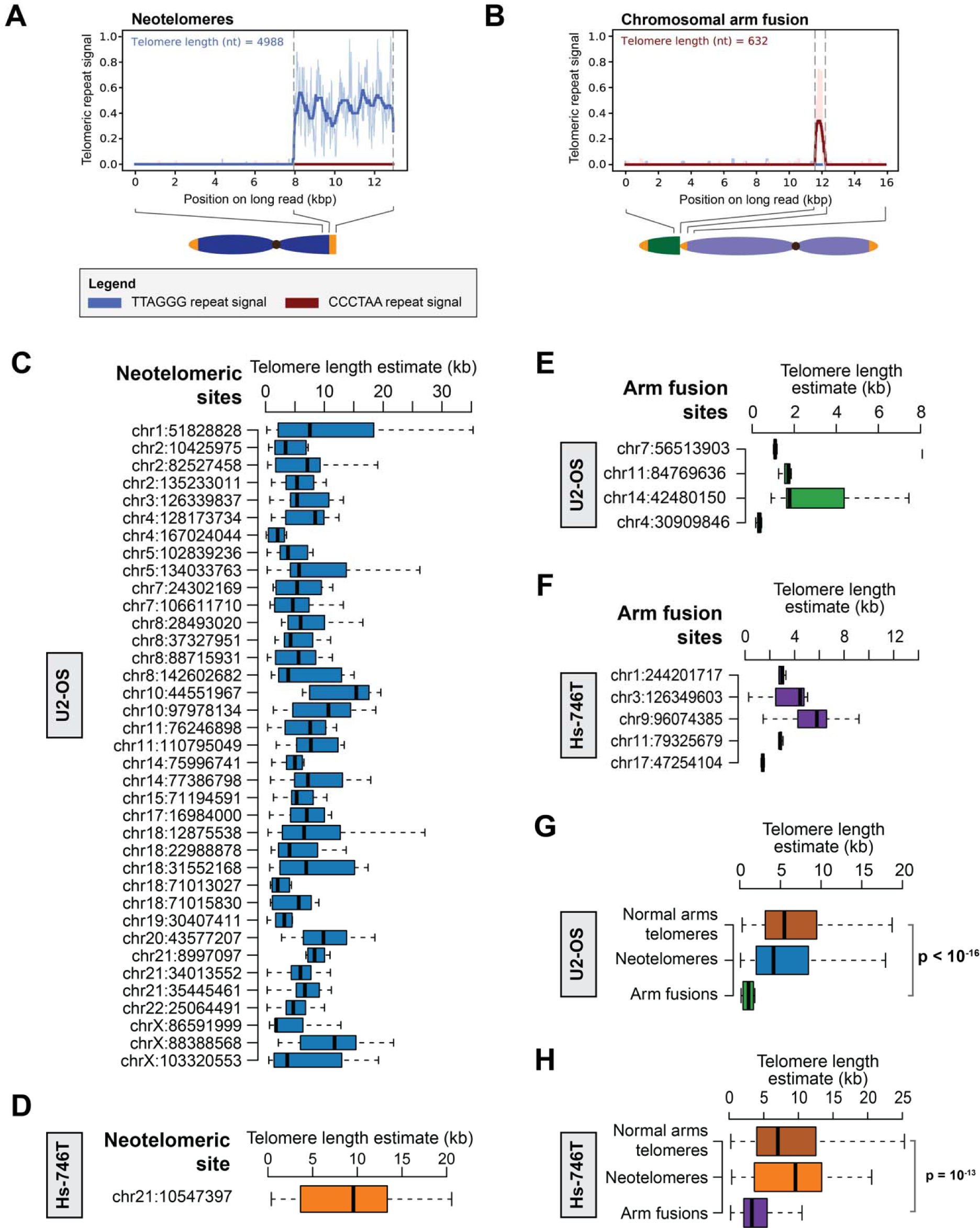
Neotelomeres have similar telomere length distribution as normal telomeres, while telomeric repeats at sites with chromosomal arm fusions are short. **(A-B)** Telomeric repeat signal observed at a representative Nanopore read with **(A)** a neotelomere in U2-OS DNA at chrX:103,320,553, and **(B)** a chromosomal arm fusion event in U2-OS DNA at chr4:30,909,846. The length of telomeric repeats on each long-read was estimated from these telomeric repeat signal profiles. Boxplots depicting the distribution of telomere length found at each neotelomere assessed by Nanopore sequencing for the **(C)** U2-OS and **(D)** Hs-746T cell lines. Boxplot depicting length of telomeric repeats assessed using Nanopore sequencing for each chromosomal arm fusion event in the **(E)** U2-OS and **(F)** Hs-746T cell lines. Note: telomere length for neotelomeres and normal chromosomal arms were only estimated using long-reads reads that start or end in telomeric repeats, while length of telomeric repeats at chromosomal arm fusions were estimated using long-reads with telomeric repeats in the middle of the read. Aggregated telomeric length of all long-reads at the normal chromosomal arms (p- and q-arms), neotelomeres, and chromosomal arm fusion events in the **(G)** U2-OS and **(H)** Hs-746T cell lines. P-values indicated in the plots were calculated using the two-sided Wilcoxon Rank Sum test. See also Figure S9.

By composite analysis of data corresponding to each class of events, we see that structurally unaltered normal chromosomal ends (p- and q-arms) have similar median telomere length (∼5kb) and similar length distribution to neotelomeres (**Figure 4G-H, Figure S9E-F)** in both the U2-OS and Hs-746T cancer cell lines. Conversely, telomeric repeats at chromosomal arm fusions are significantly shorter as compared to the other classes of events (**Figure 4G-H, Figure S9E-F)**. Together, these results indicate that neotelomeres have similar telomere length as natural telomeres and are thus possibly functional. Our results also suggest that chromosomal arms with short telomeres are more likely to undergo telomere-spanning chromosomal arm fusion events

### Somatically altered ectopic telomere repeat sequences in lung adenocarcinoma genomes

Given the results of long read analysis that demonstrated both neotelomere events, corresponding largely to the standard telomeric repeat orientation, and telomere-spanning chromosome fusion events, corresponding largely to the inverted telomeric repeat orientation, in cancer cell lines, we sought to determine whether similar events could be observed as somatic genome alterations in primary human cancers. We applied TelFuse to 95 pairs of lung adenocarcinoma tumor/normal genome sequences from The Cancer Genome Atlas, or TCGA (TCGA-LUAD) **(Table S8)**. This analysis identified 34 sites with ectopic telomere sequences in the standard orientation, and 46 sites with ectopic telomere sequences in the inverted orientation **(Tables S9 and S10)**. Putative sites of ectopic telomeric repeat sequences could be seen across the genome on almost all chromosome arms, without a particular distribution in the genome at this resolution of sample number and events (**Figure 5A**). These ectopic telomere sequences, in both the standard and inverted orientations, could be in either the centromeric or counter-centromeric direction (**Figure 5A**).

**Figure 5.**
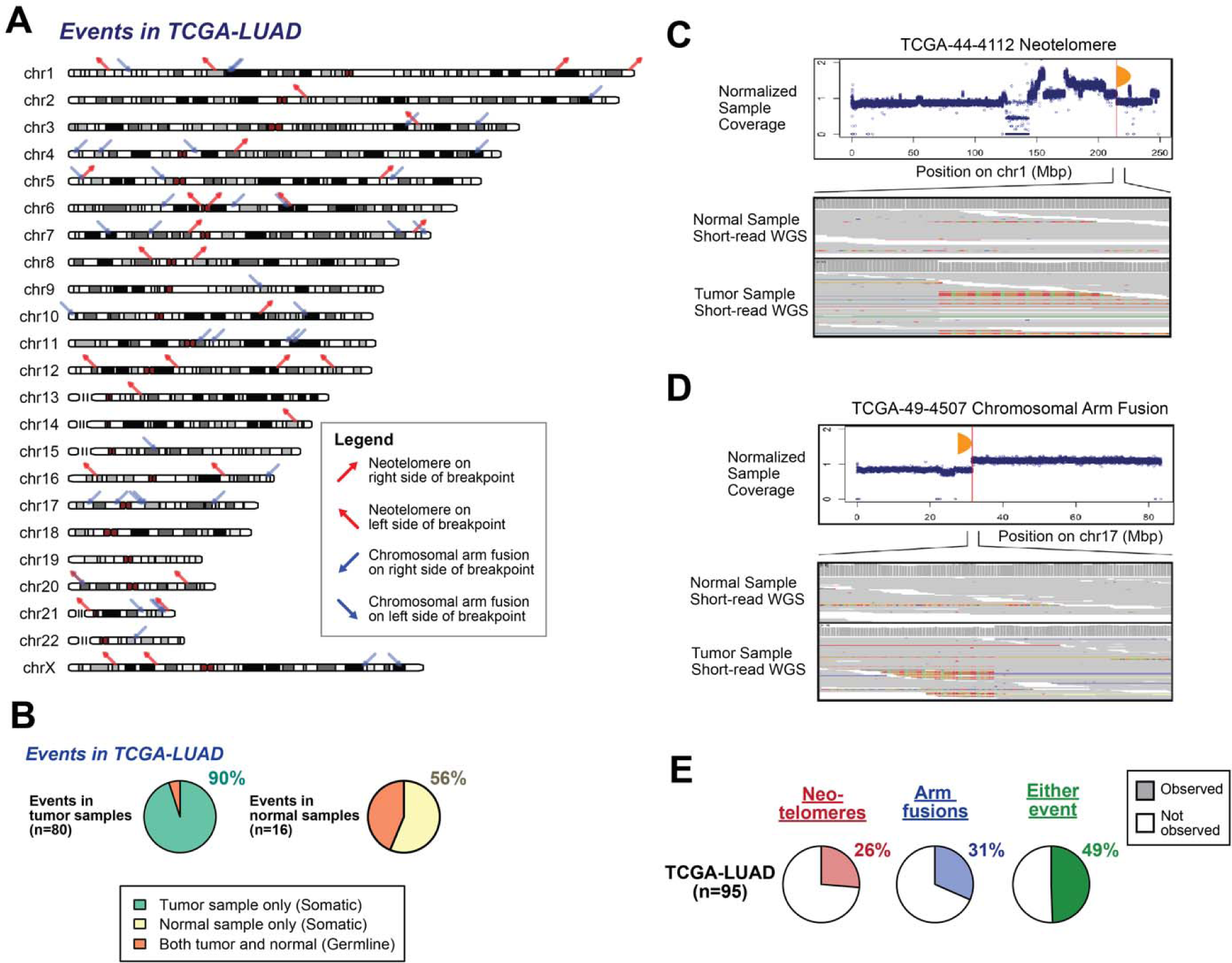
Putative neotelomeres and chromosomal arm fusion events are detected as somatic alterations in primary lung adenocarcinoma genomes. **(A)** Genome-wide distribution of putative neotelomeres and chromosomal arm fusion events in lung adenocarcinoma patient samples from The Cancer Genome Atlas (TCGA) (n=95). Neotelomeres were inferred from ectopic telomeric sequences in the standard orientation, while chromosomal arm fusion events were inferred from ectopic telomeric sequences in the inverted orientation, as described in Figure 1B, using short-read genome sequencing data. **(B)** Proportion of telomeric alterations (neotelomeres/arm fusions) that were found to be germline or somatic. **(C-D)** Examples of neotelomeres and chromosomal arm fusion events detected in tumor samples from patients with lung adenocarcinoma. **(C)** Neotelomere in tumor DNA from case TCGA-44-4112 at the site chr1:214,760,486. **(D)** Chromosomal arm fusion in tumor DNA from case TCGA-49- 4507 at the site chr17:31,537,163. Top panels: sequencing coverage at the sites of interest. Bottom panels: IGV screenshots corresponding to the neotelomere or chromosomal arm fusion events in the normal and tumor samples. **(E)** Frequency of neotelomeres and chromosomal arm fusion events in lung adenocarcinoma patient tumor samples in TCGA.

Among the standard orientation ectopic telomere repeats in the TCGA-LUAD sequence data, 32/34 sites were confirmed as somatic alterations and therefore as putative somatically generated neotelomeres by comparing the lung adenocarcinoma DNA sequence with the matched normal sequence. In addition, 44/46 of the inverted orientation repeats were confirmed as somatic alterations that are likely to represent telomere-spanning chromosomal arm fusions (**Figure 5B**). Together, among the set of 80 potential neotelomeres and chromosomal arm fusion events detected in the TCGA-LUAD tumor samples, we found that 72/80 (90%) events were only detected in the tumor sample (**Figure 5B, Table S9)**, suggesting that a large majority of calls made in tumor samples by TelFuse are somatic, even though no matched normal samples were assessed in our initial analysis.

We then performed a deeper inspection of these somatic ectopic telomere repeat sites that were detected in primary tumors. At the ectopic telomeric repeat site at chr1:214,760,486 in the patient TCGA-44-4112, at least 10 TTAGGG repeats could be observed in the primary tumor by short reads, coupled with a drop in sequencing coverage (**Figure 5C**), which is consistent with the presence of a neotelomeric site. At another site chr17:31,537,163 in the patient TCGA-49-4507, at least 6 inverted telomeric repeats of TTAGGG could be seen in the primary tumor sample by short-reads (**Figure 5D**), which may indicate the presence of a chromosomal arm fusion event given our observations with long-read sequencing of cancer cell lines. Notably, similar observations were also made at other sites with somatic ectopic telomeric repeat sequences that are consistent with potential neotelomeric or chromosomal arm fusion events **(Figure S10)** in primary lung adenocarcinoma samples. Together, this suggests that ectopic telomeric repeats in both the standard and inverted orientation can be readily observed in primary lung adenocarcinoma samples, and suggest that neotelomeres and chromosomal arm fusion events are similarly present in primary tumor samples.

All together, we observed ectopic telomeric repeats in the standard orientation and inverted orientation in 26% and 31% of the TCGA-LUAD cohort respectively (**Figure 5E**), which may point to the potential existence of neotelomeric events and chromosomal arm fusions in these samples respectively. Of note, as many as 49% of samples displayed either a neo-telomeric or chromosomal arm fusion signal, suggesting that these events are relatively common in primary tumor samples. Although this suggests an active mechanism for generation of telomeric events in cancers, we were unable to ascertain strong sequence signatures suggestive of specific telomere insertion mechanisms **(Figure S11)**.

### Germline variations leading to ectopic telomeric repeat insertions

Interestingly, we also observed 8 likely germline examples of ectopic telomere repeat sequence alterations across 4 different individuals in the TCGA-LUAD cohort (**Figure 5B, Table S9)**. A deeper exploration of these events was performed to assess the structure and features associated with these sites **(Figure S12)**. Two ectopic telomeric sites were found on the chr12q arm in both blood and tumor samples of TCGA-44-6778 at the sites chr12:54,480,142 and chr12:54,494,011, and were noted to contain a 14 kb deletion, coupled to an insertion of 6x CCCTAA repeat sequences **(Figure S12A)**. In both blood and tumor samples of the same individual at the sites chr12:25,085,740 and chr12:25,085,754 on chr12p, an insertion of 7x CCCTAA repeats was observed in tandem with duplication of a neighboring 14 bp region **(Figure S12B)**. A similar germline deletion event of 13 bp, coupled with the insertion of telomeric repeat sequences, was found in TCGA-62-A470 at chr4:184,711,090 (**Figure S12C**), while a duplication of 19 bp was coupled to a telomeric repeat insertion at chr6:170,186,789 in TCGA-44-5643 **(Figure S12D)**. Ectopic telomeric repeats could also be observed in TCGA-55-6987 at low allelic frequencies in both tumor and the adjacent normal sample **(Figure S12E)**, which may point to contamination of the normal sample or to somatic mosaicism. Together, these results indicate that ectopic telomeric repeats might be frequent germline variants, perhaps as a result of DNA repair in the presence of active telomerase^41^.

### Neotelomeres and chromosomal arm fusion events disrupt protein coding genes and are highly prevalent in cancer cell lines

In addition to allowing chromosome fusions to occur and capping truncated chromosomes, insertion of telomeric DNA might also disrupt genes, including tumor suppressors, leading to associated functional impact. To assess this possibility, we evaluated ectopic telomere sites in this study for overlap with protein coding genes. Among sites that we detected, 47% (112/240) and 47% (34/72) of sites were found to colocalize to a protein coding gene in cancer cell lines and primary lung adenocarcinomas respectively **(Table S2 and S9)**.

Notable genes with insertion events include *PTPN2*, a gene related to immunotherapy response^48^, where a neotelomere was found within the first intron, leading to a corresponding loss of the first exon and the promoter region (**Figure 6A**). Chromosomal fusion events were also found to disrupt genes, including events that led to the loss of more than half of the 5’ region of the *KLF15* and *FOXN3* genes (**Figure 6B-C**). We also observed one complex event involving telomeric DNA wherein a short neotelomere on chr1p within the *RUNX3* gen*e* then fused to the centromere of chr22/21/14. This event caused the loss of most of the gene (**Figure 6D**). Gene disruption events were also observed by long-read genome sequencing in the *NRDC* and *TENM4* genes in cancer cell lines **(Figure S13A-B)**. Interestingly, the *PTPN2*, *NRDC*, *FONX3*, and *RUNX3* genes identified in our study have putative functional roles in cancer, suggesting that the disruption of protein coding genes by neotelomeres and chromosomal arm fusions may contribute to tumorigenesis. Thus, our results indicate that neotelomeres and chromosomal arm fusion may represent an important but poorly appreciated mechanism for gene disruption.

**Figure 6.**
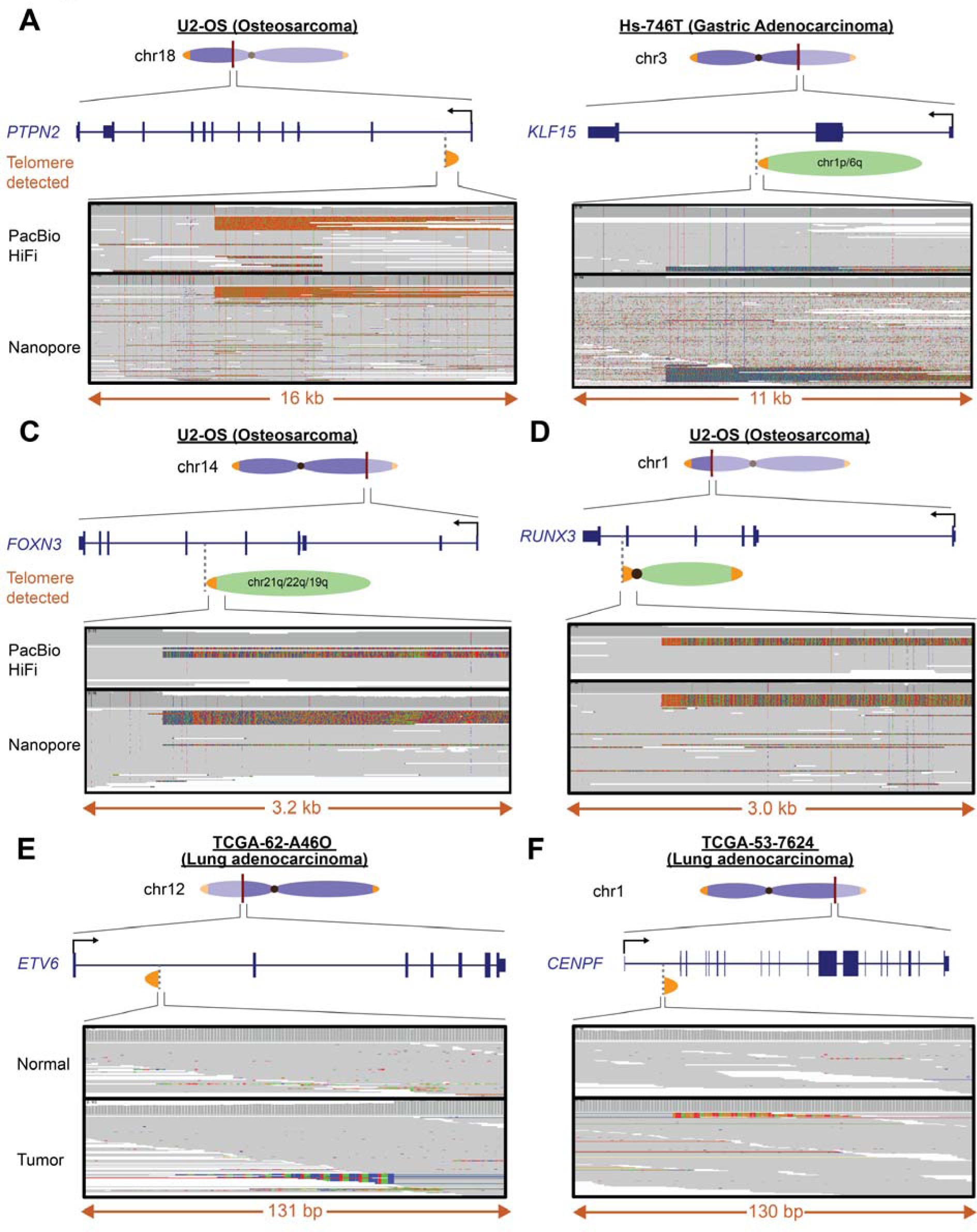
Neotelomeres and chromosomal arm fusion events disrupt protein coding genes in cancer cell lines and patient samples. **(A)** Disruption of the *PTPN2* gene in the U2-OS osteosarcoma cell line at chr18:12,875,538 with addition of a neotelomere. **(B)** Disruption of the *KLF15* gene in the Hs-746T gastric adenocarcinoma cell line associated with a chromosomal arm fusion event at chr3:126,349,603. **(C)** A chromosomal arm fusion event in the U2-OS cell line between a broken chromosome 14 and the telomere arm of chromosome 21q/22q/19q associated with disruption of the *FOXN3* gene at chr14:89,300,563. **(D)** A neotelomere in the U2-OS cell line coupled to fusion to a centromere leads to disruption of the *RUNX3* gene at chr1:24,906,321. **(E)** A putative neotelomere associated with disruption of the *ETV6* gene in a lung adenocarcinoma tumor sample derived from the patient TCGA-62-A46O at the site chr12:11,696,012. **(F)** A putative neotelomere associated with disruption of the *CEPF* gene in a lung adenocarcinoma tumor sample derived from the patient TCGA-53-7624 at the site chr1:214,609,478. See also Figure S13.

We next assessed if these gene disruption events from telomeric insertion can also be observed in primary tumor samples. In the lung adenocarcinoma sample TCGA- 62-A46O, a putative neotelomere could be observed using short-read data within the gene encoding the ETS family transcription factor, *ETV6* which is known to be associated with leukemia and congenital fibrosarcoma^47,49,50^ (**Figure 6E**). Another putative neotelomere event was observed within the gene encoding centromere protein F, *CENPF* which is thought to play a role in chromosome segregation during mitosis^51–53^ (**Figure 6F**). Putative neotelomeres and chromosomal arm fusion events were also found within the protein arginine methyltransferase gene, *PRMT7*, and the forkhead box transcription factor, *FOXP4*, genes respectively **(Figure S13C-D)**. Of note, due to the size and scale at which these neotelomeres and chromosomal arm fusion events occur, they are likely to fully disrupt these genes. Therefore, our results indicate that the formation of neotelomeres and telomere-spanning chromosomal arm fusions may represent a mechanism for gene disruption, in addition to their roles in defining gross chromosomal structure.

## Discussion

While alterations in telomere sequences are key events in cancer genome evolution, the precise nucleotide-level structure of these alterations has been hitherto inaccessible because of the inability of short-read sequence data to resolve longer repetitive sequences. Here, using long-read sequencing technologies, we delineated four types of alterations in telomere repeat sequences. First, we provide evidence that cancer cell line and primary cancer genomes contain long (several kilobase) additions of telomere repeat sequences to intra-chromosomal sites, in the standard telomere orientation (**Figure 7A**). Second, we identify telomeric repeat sequences of varying length that bridge the end of one chromosome to an intra-chromosomal site on a different chromosome (**Figure 7B**). These telomeric repeats are consistent with karyotyping analyses that have observed the attachment of chromosomal fragments to the ends of existing chromosomes^54–59^, which are key events in cancer genome evolution. Third, we observe more complex alterations where the formation of a neotelomere is followed by the fusion of the neotelomere to a second intra-chromosomal location (**Figure 7C**). Fourth, we observe fusions that link centromeric to telomeric sequence repeats (**Figure 7D**). The implications of several of these alterations are described below.

**Figure 7.**
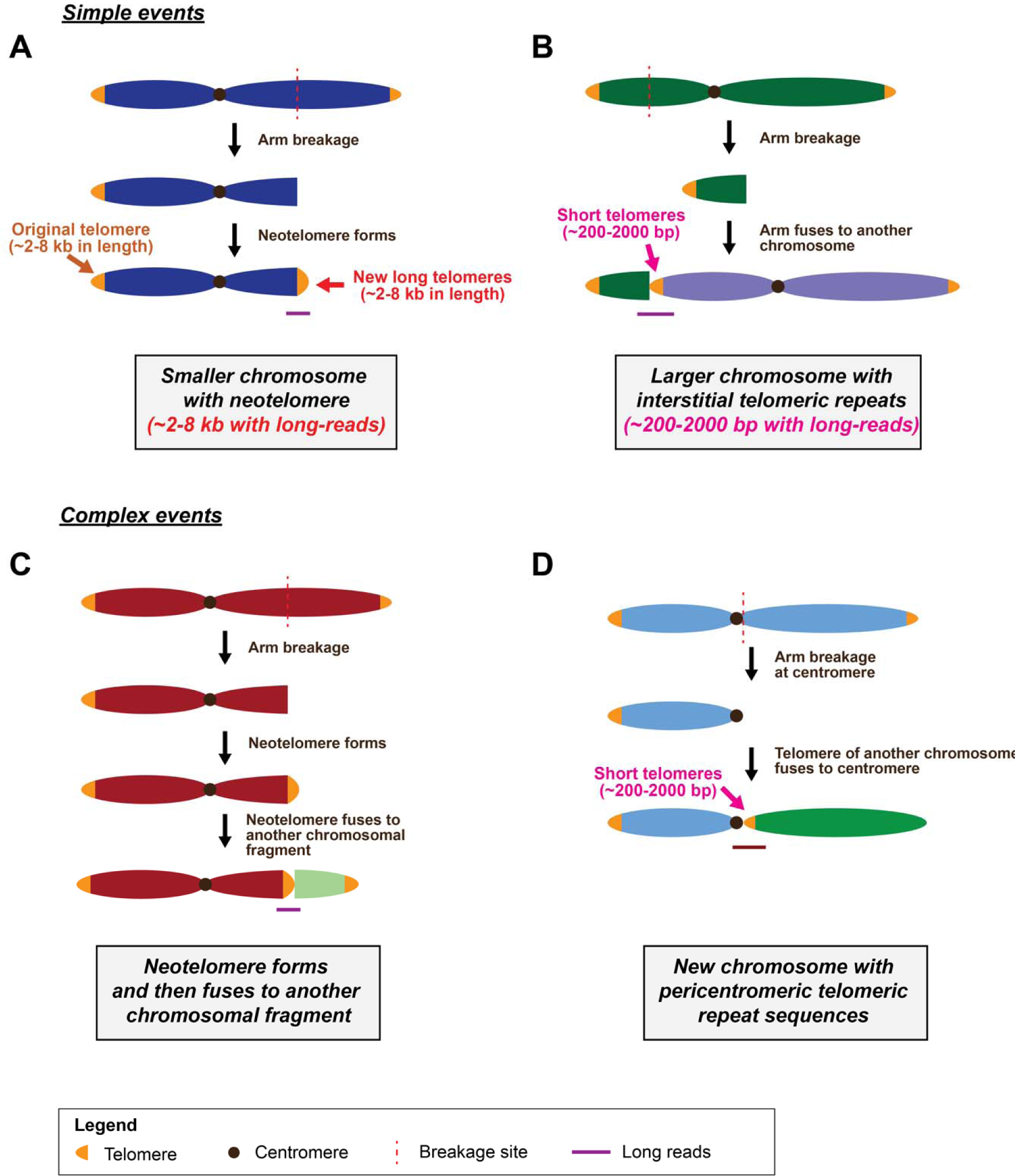
Possible models that can account for the different types of telomeric repeat sequences observed in this study. **(A)** A neotelomere can form after a chromosomal arm breakage event. This leads to the generation of a smaller chromosome with a neotelomere, similar in repeat length to telomeres found on a normal chromosomal arm. **(B)** Chromosome arm fusion where a broken chromosomal arm can fuse to another chromosome with very short telomeres. This generates a larger chromosome with interstitial telomeric repeat sequences in the middle of the chromosome**. (C)** Complex alteration where neotelomere formation is followed by the fusion of this neotelomere to another chromosomal fragment. This leads to the observation of long-reads in our study which contains telomeric repeat sequences, flanked on both sides by intra-chromosomal sequences. **(D)** A complex telomeric alteration involving a chromosomal arm break at or very near to the centromere, which is fused to another chromosomal arm with very short telomeres. The resultant new chromosome has pericentromeric telomeric repeat sequences. Purple line: parts of the model supported by long-read genome sequencing data.

A previous study, analyzing short-read genome sequencing of patients’ cancer samples from the Pan-Cancer Analysis of Whole Genomes (PCAWG) project, was able to identify a number of intra-chromosomal telomeric repeat insertion sites^23^. In comparison to this previous study, our work shows that telomere length at these repeat insertion sites can be estimated using long-read sequencing, and the underlying sequence structure can be analyzed in the context of adjacent sequences compared to free telomeric ends. This technical advance allowed us to differentiate intra-chromosomal telomeric repeat sites based on the orientation of the telomeric repeat sequences. By integrative analysis of long-read genome sequencing, spectral karyotyping, coverage analysis, and short-read genome sequencing, we demonstrated the existence of multi-kilo base-pair long neotelomeres at sites of putative chromosomal arm breakages, corresponding to telomeric repeats in the standard orientation. We further provided evidence for the presence of these standard orientation telomere repeats representing neotelomeres in primary tumor sequence data of lung adenocarcinoma (LUAD) from TCGA. Further, powered by long-read genome sequencing, we were able to reliably show that sites with inverted telomeric repeat sequences represent fusion of chromosomal arms spanning short telomere sequence repeats, also found in TCGA LUAD data. Together, our study provides support for the existence of neotelomeres and chromosomal arm fusion events in cancer genomes, and also provides insights into the cause of their occurrence.

A recent experimental study generated double-strand breaks in cells over-expressing telomerase, leading to the addition of neotelomeres at a subset of these breaks^41^. Our study provides genomic evidence for a signature of neotelomere addition in cancer cell lines and cancer genomes, complementary to this experimental evidence. The location and the unbounded structure of these repeats suggest that they are likely to be functional neotelomeres. Taken together, the cellular experiments and genomic observations support a model where neotelomere addition by telomerase, nucleating at sites of double strand breaks, can be a common step in tumorigenesis.

The generation of new chromosomes via chromosomal rearrangements is a key element of cancer genome evolution and also occurs during the course of evolution and speciation^60–62^. Some of our findings using long-read sequencing of cancer genomes mirror long-standing observations in genomes of many organisms. Interstitial telomeric repeats have been identified in the genomes of many vertebrates, including primates and the pygmy tree shrew^63–65^, akin to those found at sites of chromosomal arm fusions in cancer cell lines (**Figure 7B**). Furthermore, interstitial telomeric sequences have been observed close to centromeres in the genomes of diverse organisms including Chinese hamster, Arabidopsis, and the European grayling^65–67^. These structures, termed pericentromeric telomeric repeats, were similarly observed by long-read genome sequencing in the U2-OS cancer cell line in our study (**Figure 7D**). Overall, the study of telomere repeat alterations also provide an understanding into how new chromosomes originate during the course of evolution and speciation, as well as during cancer genome evolution.

Looking at the genome beyond telomeric repeats, repetitive elements constitute approximately half the human genome^13–15^. However, we have not yet been able to understand genome structure and alterations at a detailed level because of the technical limitations of short-read sequencing, which is unable to span or completely delineate the precise structure of these repeat elements. Here, using telomeres as a salient example, we show how long-read genome sequencing can be used to drive discoveries of functional importance in highly repetitive regions of the cancer genome, and also inform the analysis of existing short-read data. As a bridge to a future where universal long-read sequencing is technically and economically feasible, our study provides a framework to assess short-read genome sequencing data for genome alterations within highly repetitive regions, that can be followed by long-read sequencing and complete analysis of selected samples. Significantly, given that >95% of repetitive sequences in the genome are estimated to be <8 kb in length^15^, long-read sequencing data that is typically generated at >10 kb in length **(Figure S3)** would enable the majority of previously neglected alterations in the cancer genome to be completely resolved. Thus, our study highlights the utility of long-read genome sequencing in the study of chromosomal scale structures in cancer and beyond. This analysis may have functional implications as we observed the disruption of protein coding genes by neotelomeres and chromosomal arm fusions. More broadly, the identification of these gene disruptions points to the potential role that other repetitive elements may play in gene disruption as well as activation events and to the discovery opportunity provided by long-read cancer genome sequencing.

There are a few limitations associated with our study. First, in contrast to a recent yeast genomic study in which the end of each telomere was tagged^68^, it is difficult to assess if telomeric repeats containing long-reads analyzed in our study captured the telomeres end-to-end. As such, telomere length estimates made in our study may underestimate the true length of telomeres. Further, it also known that the sub-telomeres at normal chromosomal arms contain telomere-like sequences and short internal telomeric repeats close to long stretches of perfect (TTAGGG)_n_ repeats^44,69^. However, it is unclear if these sequences should be included in the computation of telomere length estimates performed in our study.

In summary, we have used long-read sequencing to demonstrate the generation of neotelomeres, and of chromosome arm fusions that span telomere repeats, in human cancer cell lines and then provided evidence for these alterations in primary human lung adenocarcinoma genomes. This study provides detailed insight into the process of telomere maintenance in human cancer. Further long-read sequencing studies of cancer genomes could help to elucidate the potential role of somatic alterations in highly repetitive regions of the human genome in cancer pathogenesis. More broadly, long-read sequencing analyses may also provide insights into chromosomal rearrangements that drive genetic diseases and evolution.

## Acknowledgements

We thank all members of the Matthew Meyerson and Heng Li labs for helpful comments and inputs on the work. We would also thank Jidong Shan (Albert Einstein College of Medicine) for generating spectral karyotyping results, and for inputs on the analysis of cytogenetics data. We further thank Jodi Hirschman for assistance with edits to our manuscript.

## Author contributions

K.T.T. and M.M. initiated the study of telomeres in cancer with long-read genome sequencing. K.T.T developed computational methods and designed computational analyses with input from H.L. and M.M. K.T.T. performed most computational analyses in this study. M.G.J. assisted with computational analysis of TCGA-LUAD dataset. M.S. generated DNA samples of cancer cell lines used for long-read genome sequencing, and performed an initial long-read sequencing run. K.T.T. wrote the initial draft of the manuscript with input from M.M., M.L.L, and H.L. M.M. and H.L. jointly supervised the work. All authors read, revised, and approved the submission of the manuscript.

## STAR★Methods

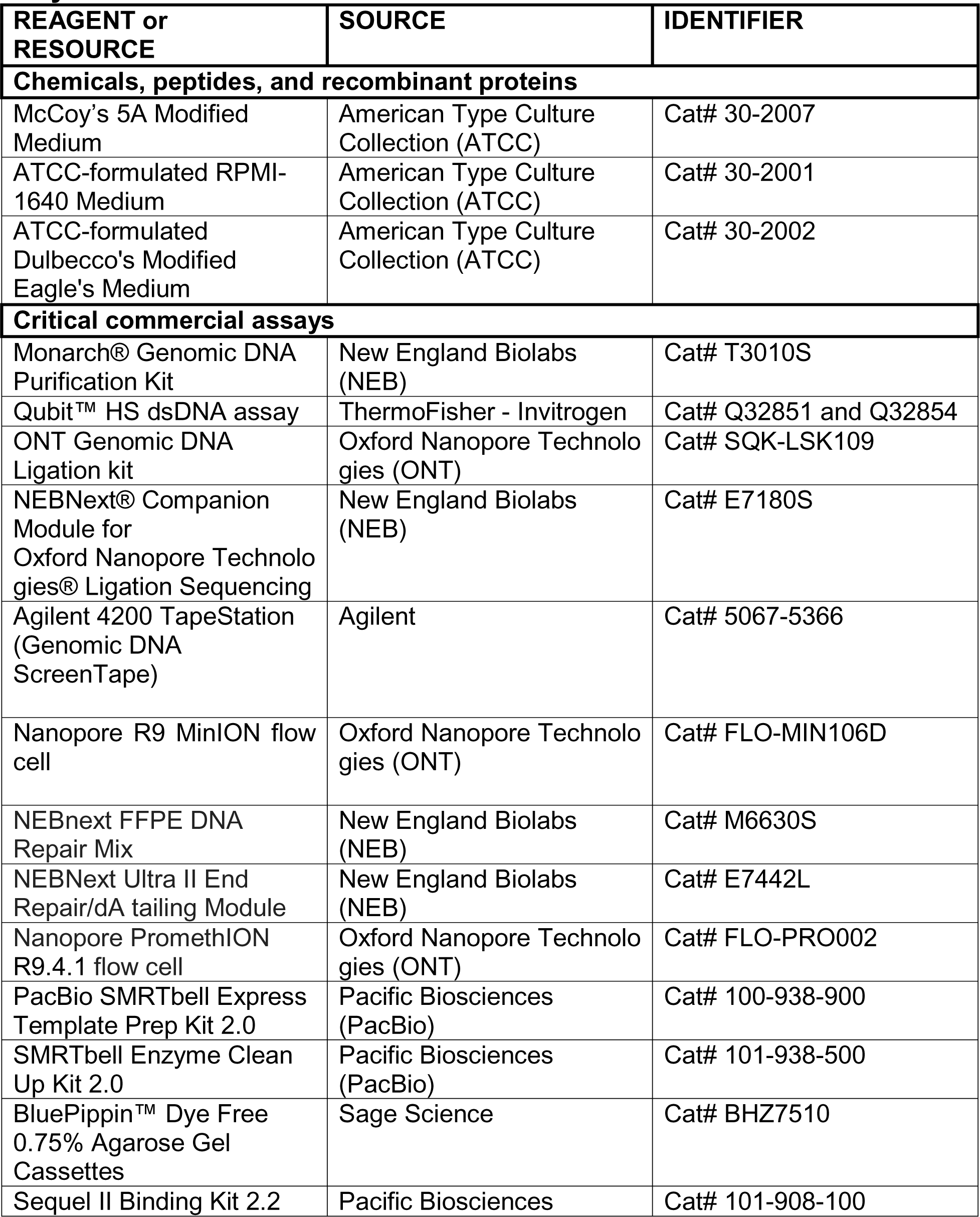
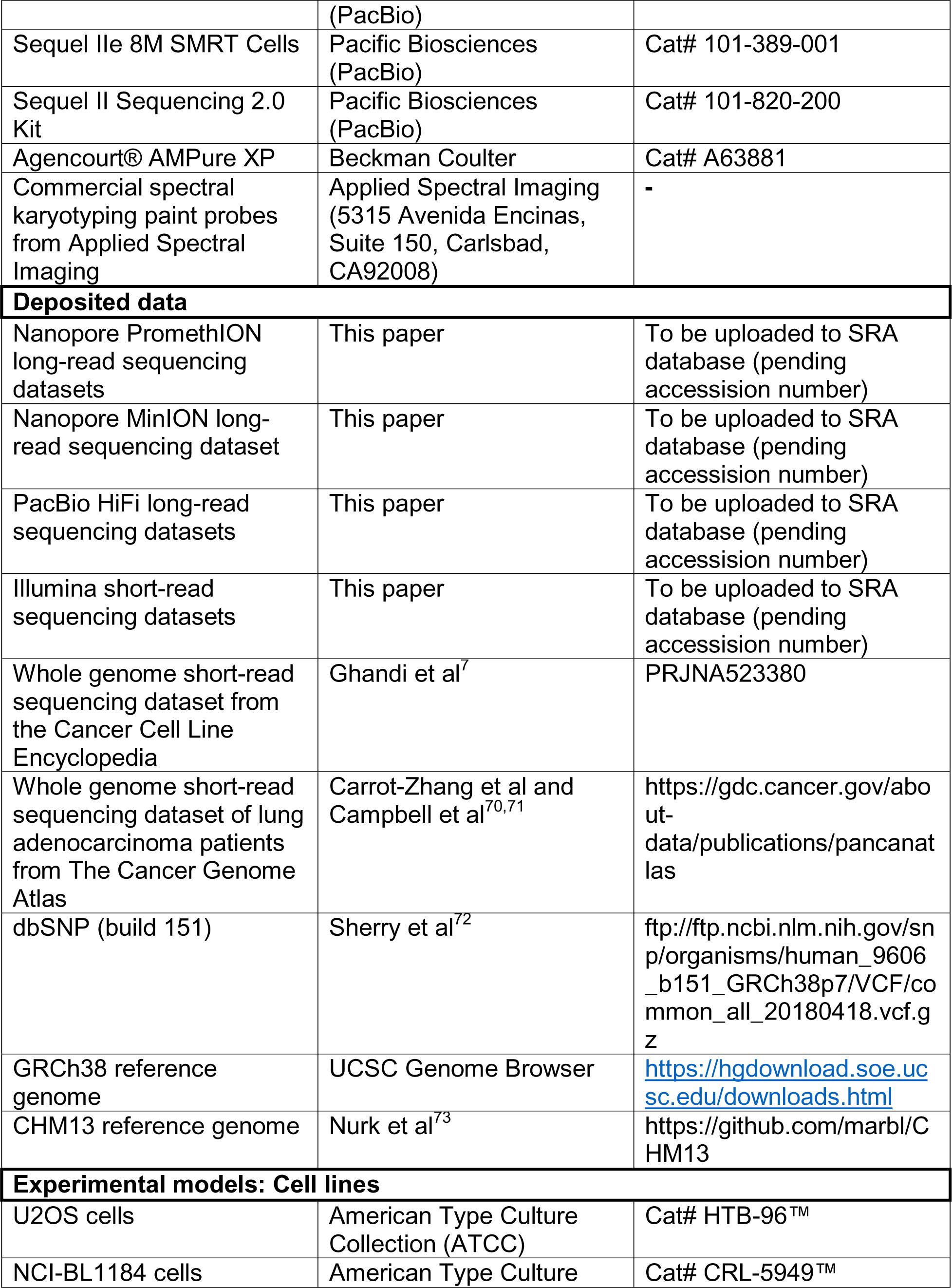
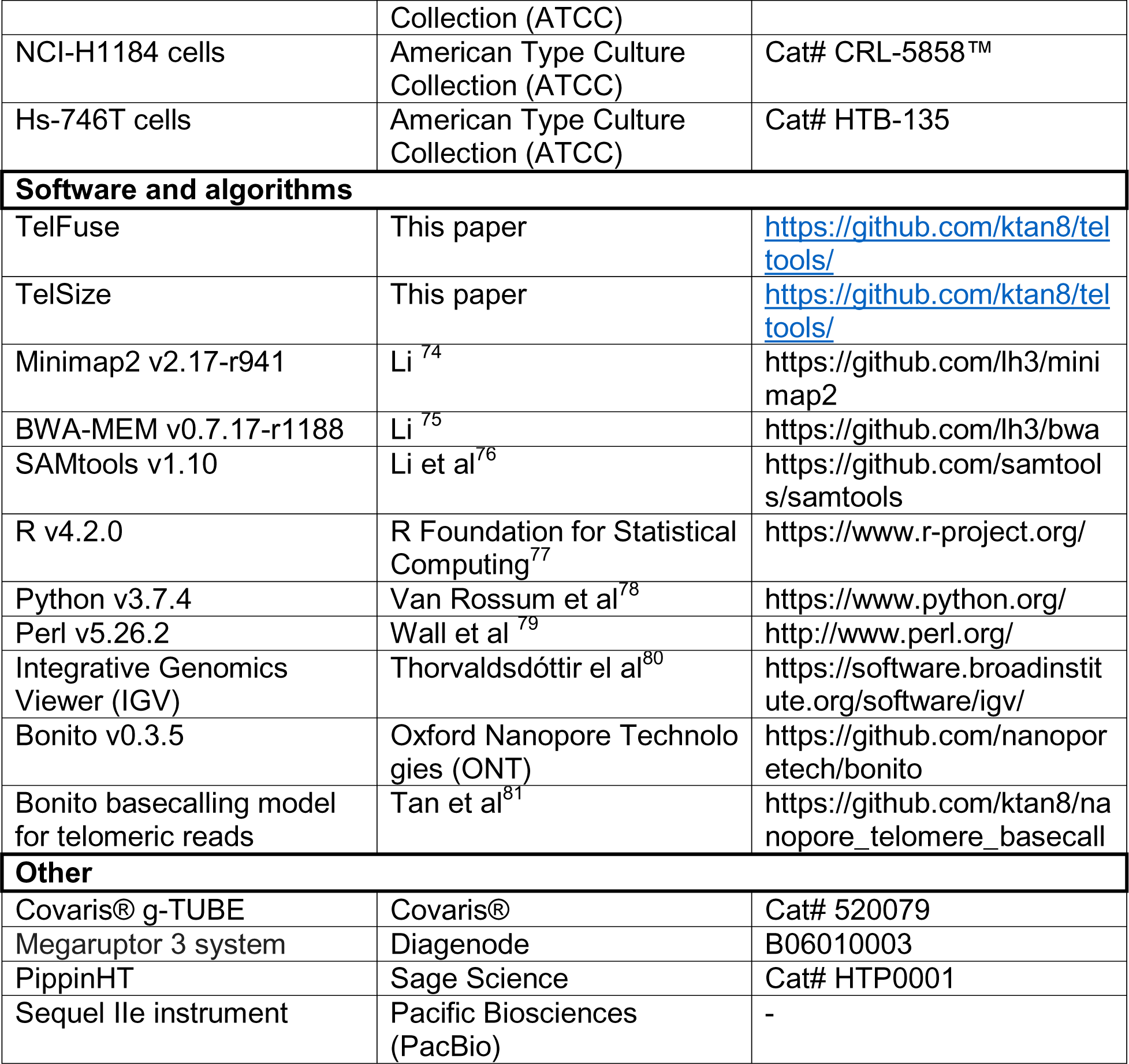
Key resources table.

## Resource availability

### Lead contact

Further information and requests for resources should be directed to and will be fulfilled by the lead contact, Matthew Meyerson.

### Materials availability

This study did not generate new unique reagents.

## Methods details

### CCLE whole genome sequencing dataset

CCLE dataset^7^ was downloaded from the European Nucleotide Archive under the study accession number (PRJNA523380). Specifically, only whole genome sequencing (WGS) datasets from the study was obtained. A full list of accession numbers corresponding to the CCLE WGS dataset used in this study is indicated in Table S1.

### Lung adenocarcinoma whole genome sequencing dataset

Whole genome short-read sequencing dataset of lung adenocarcinoma patients^70,71^ from The Cancer Genome Atlas were downloaded from the GDC Data Portal (https://portal.gdc.cancer.gov/). The list of accession numbers corresponding to samples analyzed for this study is indicated in Table S8.

### Identification of candidate new telomeres and chromosomal arm fusion events from short reads

Candidate short read pairs with at least two consecutive telomeric repeat sequences (TTAGGG)_2_ in either reads in the pair were first extracted to narrow down the number of read pairs for subsequent analysis. Specifically, this was done by applying a custom Python script in the TelFuse package to each whole genome sequencing dataset.

Candidate read pairs were then remapped to the reference genome (GRCh38) with BWA-MEM (version 0.7.17-r1188)^75^ with default parameters. A custom Python script in the TelFuse package was then used to extract all sites with soft-clipped regions on the mapped reads. Soft-clipped sequences from all reads at each unique genomic site was then used to generate a consensus sequence. A corresponding average sequence identity of the soft-clipped sequences to the consensus was also calculated.

To then filter this list of candidate sites for potential new telomeres and chromosomal arm fusion events, a series of filters were applied. Specifically, we ensured that (i) each site is supported by at least 3 reads, (ii) has an average sequence identity to the consensus of ≥ 95%, (iii) average mapping quality ≥30, (iv) found more than 500kb from each end of the chromosome as defined by the reference genome, (v) is not found in more than one sample in the “panel of normal” constructed from these samples, and (vi) contains the circular permutations of (TTAGGG)_2_ or (CCCTAA)_2_ sequence in the soft-clipped sequences immediately after the breakpoint.

The candidate sites were then further subdivided into sites with telomeric repeats in the standard or inverted orientation, depending on the orientation of telomeric repeat sequences with respect to the genomic loci of interest.

### Cell culture

U2-OS cells (ATCC® HTB-96™) were cultured in McCoy’s 5A Medium Modified (ATCC cat no. 30-2007) with 10% FBS. Cell lines NCI-BL1184 (ATCC cat no. CRL-5949™) and NCI-H1184 (ATCC cat no. CRL-5858™) were cultured in ATCC-formulated RPMI-1640 Medium (ATCC cat no. 30-2001) supplemented with FBS at 10%. Hs-746T cells (ATCC cat no. HTB-135) were cultured in ATCC-formulated Dulbecco’s Modified Eagle’s Medium (ATCC cat no. 30-2002) supplemented with 10% FBS

### High molecular weight DNA extraction

High molecular weight (HMW) DNA was isolated using a Monarch® Genomic DNA Purification Kit (NEB, cat no. T3010S). DNA was quantified with a Qubit™ HS dsDNA assay (ThermoFisher, cat no. Q32851) followed by verification of HMW DNA integrity by electrophoresis on an Agilent 4200 TapeStation (Genomic DNA ScreenTape, cat no. 5067-5366).

### MinION Library Preparation

Sequencing libraries were prepared for the Oxford Nanopore Technologies (ONT) platform using the ONT Genomic DNA Ligation kit (ONT, cat no. SQK-LSK109). Briefly, HMW U2OS DNA was fragmented to ∼20 Kb using a Covaris® g-TUBE (cat no. 520079) followed by SPRI-cleanup (Agencourt® AMPure XP, Beckman Coulter, cat no. A63881). Fragmented material was quantified with a Qubit™ dsDNA HS Assay Kit (Invitrogen™, Catalog number: Q32851). One microgram of HMW U2OS DNA was end-repaired and A-tailed (NEBNext® Companion Module for Oxford Nanopore Technologies® Ligation Sequencing, cat no. E7180S) followed by adapter ligation. For sequencing 100 fmols of library material was loaded on an R9 flow cell (cat no. FLO-MIN106D).

### PromethION Library Preparation

Sequencing libraries for PromethION sequencing was prepared using the Genomic DNA by Ligation kit (SQK-LSK109) provided by Oxford Nanopore Technologies according to the recommended protocol (Version GDE_9063_v109_revT_14Aug2019) with slight modifications to the amount of input DNA used and the equipment used for shearing of the DNA. Briefly, 2.5 ug of high molecular weight genomic DNA was sheared to 20kb using a Megaruptor 3 system (Diagenode, cat no. B06010003). DNA repair and end-prep was then performed using the NEBnext FFPE DNA Repair Mix and NEBNext Ultra II End Repair/dA tailing Module reagents in accordance with the manufacturer’s instructions followed by cleanup with AMPure XP beads. Ligation of adapters was then performed using the Ligation Sequencing kit (SQK-LSK109) according to manufactuer’s instructions, followed by loading onto a PromethION R9.4.1 flowcell (Oxford Nanopore, cat no. FLO-PRO002).

### PacBio HiFi Library Preparation

For CCS library preparation, ≥3 ug of high molecular weight genomic DNA (more than 50% of fragments ≥40 kb) was sheared to ∼15 kb using the Megaruptor 3 (Diagenode B06010003), followed by DNA repair and ligation of PacBio adapters using the PacBio SMRTbell Express Template Prep Kit 2.0 (100-938-900) and removal of incomplete ligation products with the SMRTbell Enzyme Clean Up Kit 2.0 (PacBio 101-938-500). Libraries were then size-selected for 15 kb +/- 20% using the PippinHT with 0.75% agarose cassettes (Sage Science). Following quantification with the Qubit dsDNA High Sensitivity assay (Thermo Q32854), libraries were diluted to 60 pM per SMRT cell, hybridized with PacBio V5 sequencing primer, and bound with SMRT seq polymerase using Sequel II Binding Kit 2.2 (PacBio 101-908-100). CCS sequencing was performed on the Sequel iIe instrument using 8M SMRT Cells (101-389-001) and Sequel II Sequencing 2.0 Kit (101-820-200), utilizing PacBio’s adaptive loading feature with a 2 hour pre-extension time and 30 hour movie time per SMRT cell. Initial quality filtering, basecalling, adapter marking, and CCS error correction was done automatically on board the Sequel iIe.

### Base calling of Nanopore sequencing data

Base calling of Nanopore sequencing data in this study was performed using Bonito (Version 0.3.5) with the default dna_r9.4.1 basecalling model. However, the default Nanopore basecalling model leads to frequent strand-specific base calling errors at telomeric repeats in our dataset, with (TTAGGG)_n_ being miscalled as (TTAAAA)_n_, and (CCCTAA)_n_ being miscalled as (CTTCTT)_n_ and (CCCTGG)_n_, akin to what we had previously reported^81^. As such, telomeric reads was extracted using a pipeline that we had previously developed, followed by re-basecalling using a basecalling model that was previously tuned to correct these errors^81^.

### Extraction of candidate telomeric long reads for detailed analysis by TelSize

Long reads containing telomeric repeats were extracted by first enumerating the number of (TTAGGG)_2_ and (CCCTAA)_2_ motifs on each read using custom Perl scripts. Long reads containing at least four of these motifs were then defined as candidate telomeric repeats. Of note, a low cutoff was deliberately set here to more sensitively identify long-reads with telomeric repeats for detailed analysis by TelSize.

### Estimation of telomere length from noisy long reads

As the telomeric long reads generated by Nanopore sequencing was relatively noisy, the length of telomeric repeats could not be readily inferred from the reads. To address this, we scanned each telomeric long read for instances of the telomeric repeat sequence (TTAGGG), or its reverse complement (CCCTAA). A vector representing positions where each of these motifs were observed was then generated. We then applied a moving average filter with window size 50 on this profile, followed by a moving median filter with window size 501. A minimum telomeric repeat signal of ≥ 0.35 was then applied to define a region as telomeric. The size of the telomeric repeat region was then established to determine the length of telomeric repeats on the long read, the localization of these sequences on the long-reads, and if (CCCTAA)_n_ or (TTAGGG)_n_ repeats were observed.

Specifically, long-reads were classified into five different classes: full telomeric – long-reads that contains telomeric repeat sequences end-to-end, left telomeric – long-reads that contains telomeric repeat sequences on the left edge of the long-read, right telomeric – long-reads that contains telomeric repeat sequences on the right edge of the long-read, intra-telomeric – long-reads that contains telomeric repeat sequences in the middle of the single long-read, and non-telomeric – long-reads that do not contain significant telomeric repeat signal throughout the long-read. These telomeric repeat signal can also occur as either (TTAGGG)_n_ or (CCCTAA)_n_ repeats, and these information are further reported.

This package for telomeric long read extraction and estimation (telSize) is available at the following github repository (https://github.com/ktan8/teltools/).

### Analysis of telomeric repeat length at neotelomeres and chromosomal arm fusion sites

To assess length of telomeric repeats at neotelomeres and chromosomal arm fusion sites, only left telomeric, right telomeric, and intra-telomeric reads were considered. Specifically, for neotelomeric events, only reads with telomeric repeat regions found at the 5’ or 3’ end of the read (i.e. left telomeric and right telomeric reads) was considered to ensure that these reads correspond to a terminal region of a genomic locus. In the context of chromosomal arm fusion events, we require that the telomeric region be situated within the long-read (i.e. intra-telomeric reads that are flanked by non-telomeric repeats on both sides) to ensure that reads analyzed at these loci represent chromosomal arm fusion events.

For these telomeric repeat containing reads, sequences corresponding to the telomeric repeat region were trimmed off. The remaining non-telomeric sequences of each read were then mapped to the GRCh38 reference genome with minimap2 (Version 2.17- r941). Primary read mappings in the PAF format were then extracted and analyzed using custom R scripts in order to assess mapping coordinates of these sequences. For each site of interest that was identified using short-read data, telomeric repeat containing long-reads that mapped to a ±100 bp region of each site were extracted. Telomere length estimates for long-reads at each neotelomeric and chromosomal arm fusion sites were then reported as per Figure 4.

### Analysis of telomeric repeat length at normal chromosomal arms

To assess length of telomeric repeats at normal chromosomal arms, only left telomeric and right telomeric reads were considered, akin to the neotelomeric sites. Sequences corresponding to the telomeric region were similar trimmed off. The remaining non-telomeric repeat sequences were mapped to the CHM13 v2.0 reference genome using minimap2 (Version 2.17-r941) as the sub-telomeric region of this reference genome is complete in in contrast to the GRCh38 reference genome. Reads that mapped to the terminal 500kb region of each chromosomal arm were classified as telomeric reads originating from normal chromosomal arms.

### Copy number profiles

To generate copy number profiles of the cancer cell lines from the CCLE, the total sequencing coverage of each 10 kb bin was calculated using the bedcov function SAMtools (v1.10)^76^ with default parameters. The coverage was then normalized to a per-basepair level and is as depicted.

For lung adenocarcinoma samples which has a matched normal samples, the normalized sample coverage across each chromosome was calculated as follows. The sequencing coverage for each 10kb bin was calculated for both the tumor and matched normal sample using the bedcov function in SAMtools (v1.10)^76^. These values were then normalized by the total read count of each dataset, and the ratio between the tumor and normal sample calculated to obtain the normalized sample coverage.

### Analysis of BAF

As no matched normal samples were sequenced for each of the cancer cell lines, heterozygous germline variants cannot be directly assessed and used in the generation of allelic ratio plots. Allelic ratios was thus assessed using a set of common germline SNPs from the dbSNP database (GRCh38.p7 build 151)^72^ (ftp://ftp.ncbi.nlm.nih.gov/snp/organisms/human_9606_b151_GRCh38p7/VCF/common_all_20180418.vcf.gz). Specifically, the list of common SNPs are defined by the dbSNP database as SNPs that are found with a minor allele frequency of at least 0.01 in the 1000 genomes project.

Custom Python scripts and SAMtools mpileup (v1.10) were then used to enumerate all four possible bases at each SNP site (base quality ≥ 20). The allelic ratio was then calculated as the ratio of the variant base (as defined by dbSNP) count versus the sum of the reference and variant base count. Only sites with a coverage of at least 15x were plotted.

Sequence signatures at sites with new telomeres and chromosomal arm fusion events.

The sequence signature at each new telomeric and chromosomal arm fusion site was analyzed using the consensus soft-clipped sequences identified by TelTools, and the sequence extracted from the reference genome at each site. The sequence signature at each new telomere and chromosomal arm fusion was then analyzed by (i) identifying the frequency of each telomeric 6-mer in each soft-clipped sequence, and by (ii) assessing the sequence motif of the telomeric region and genomic region.

### Spectral Karyotyping

DNA Spectral Karyotyping Hybridization was performed according to the protocol of commercial spectral karyotyping paint probes from Applied Spectral Imaging (5315 Avenida Encinas, Suite 150, Carlsbad, CA92008). Briefly, the slides were dropped in Thermotron and aged for 3-5 days in a 37°C oven. The slides were then checked under the microscope before hybridization. A series of four steps were then performed on these slides to generate the spectro karyotype of the cell lines:

### (1) Trypsin Treatment

The slides were washed briefly in Earl’s medium, and then treated with Trypsin/EDTA solution. Washing was then performed in water and then dehydrated in ethanol series of 70%, 80% and 100% for 2 minutes each followed by air-dying of the slides.

### (2) Chromosome Denaturation

The slides were treated in 2XSSC buffer for 2 minutes and then dehydrated in Ethanol series for 2 minutes each. Denaturation of the slides was then performed at 72°C in denaturation solution for 1.5 minutes. This is followed immediately by placing the slides in cold ethanol series to dehydrate the slides, and then air drying.

### (3) Probe Denaturation and hybridization

The probe was denatured by incubating the probe at 80°C in a water bath for 7 minutes. The denatured Spectral Karyotyping reagent was then applied to the denaturized chromosome preparation and incubated at 37°C for 5-6 days.

### (4) Detection, imaging and karyotyping

The slides were washed in 0.4XSSC at 72°C for 2 minutes and then dipped in 4XSSC/Tween-20 for 1 minutes. Cy5 staining reagent was then applied and incubated at 37°C for 40 minutes.

The slides were then washed 3 times in washing solution, and then mounted with anti-fade DAPI. After which, the slides are ready for spectral imaging. Rearrangements were defined with nomenclature rules from international Committee in Standard Genetic Nomenclature for Human.

## Data and code availability

TelFuse and TelSize developed for this study are available at https://github.com/ktan8/teltools/. Long-read genome sequencing data generated for this study would be deposited in the SRA database prior to the publication of the manuscript.

## Supplemental Information

### Supplementary Figure Legends

**Figure S1.**
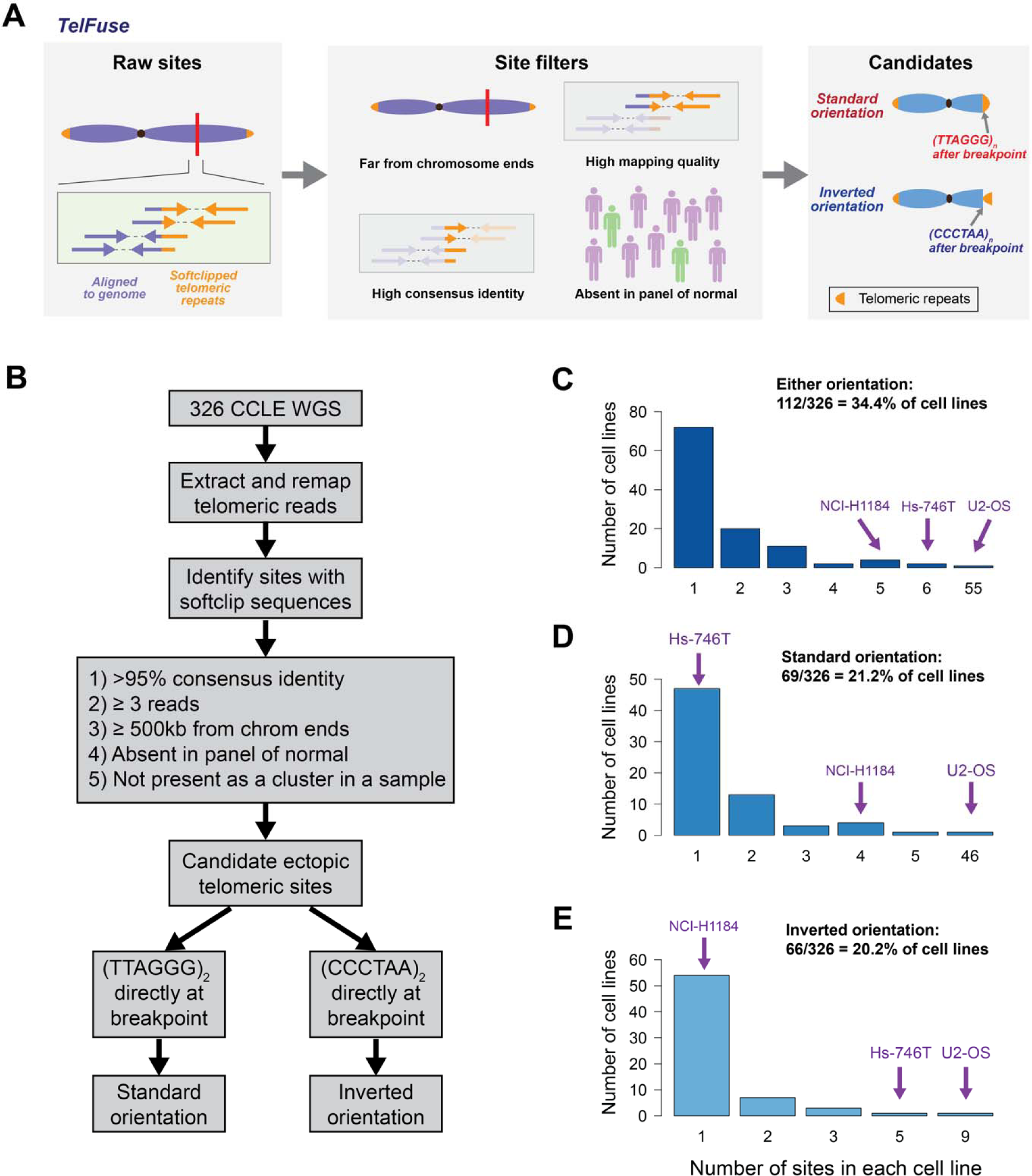
Overview of the TelFuse methodology and the distribution of ectopic telomeric events in cancer cell line genomes. **(A)** Overview of the TelFuse computational method to identify ectopic telomere repeat sequences with short-reads genome sequencing. Raw sites were first identified from partially mapped read-pairs which also contain telomeric repeats on unmapped portions of these reads. A stringent set of filters was then applied to ensure the specificity of these calls. These candidate sites were then classified by the orientation of telomeric repeats after the breakpoint. Specifically, sites with (TTAGGG)_n_ repeats after the breakpoint have telomeric repeats in the standard orientation, akin to a standard telomere. Conversely, sites containing (CCCTAA)_n_ telomeric repeats after the breakpoint have telomeric repeat sequences in the inverted orientation. **(B)** Detailed overview of method used for the identification of ectopic telomeric repeat sites. (C-E) Number of ectopic telomeric repeat sites found in the genome of each cancer cell line (n=326 cell lines) in **(C)** either orientation, in **(D)** the standard orientation, and in (E) the inverted orientation. The three cell lines (U2-OS, Hs- 746T, and NCI-H1184) which have a high frequency of ectopic telomeric events, and which were selected for long-read genome sequencing are as indicated.

**Figure S2.**
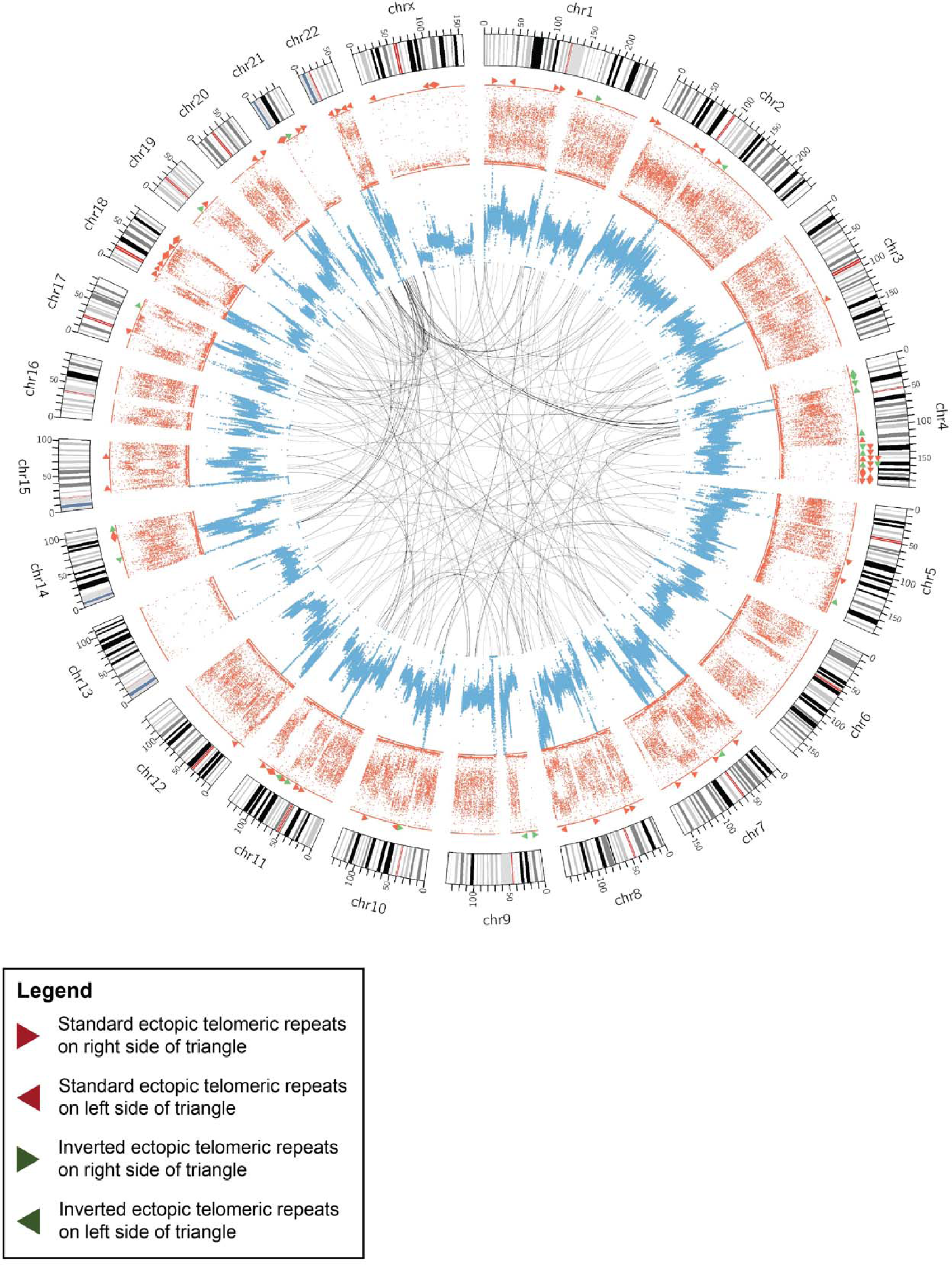
Circos plot depicting locations of ectopic telomeric repeat sequences identified in the U2-OS cell line by TelFuse. Locations of ectopic telomeric repeats in the standard orientation are labelled with red triangles, while those in the inverted orientation are labelled with green triangles. Triangles pointing in the anti-clockwise direction indicates that the ectopic telomeric repeat sequences are found on the anti-clockwise edge, while triangles pointing in the clockwise directions indicates that ectopic telomeric repeat sequences are on the clockwise edge. The allelic ratios are depicted in red, while the sequencing coverage is labelled in blue. The inner most circle depicts translocation events detected in the cell line.

**Figure S3.**
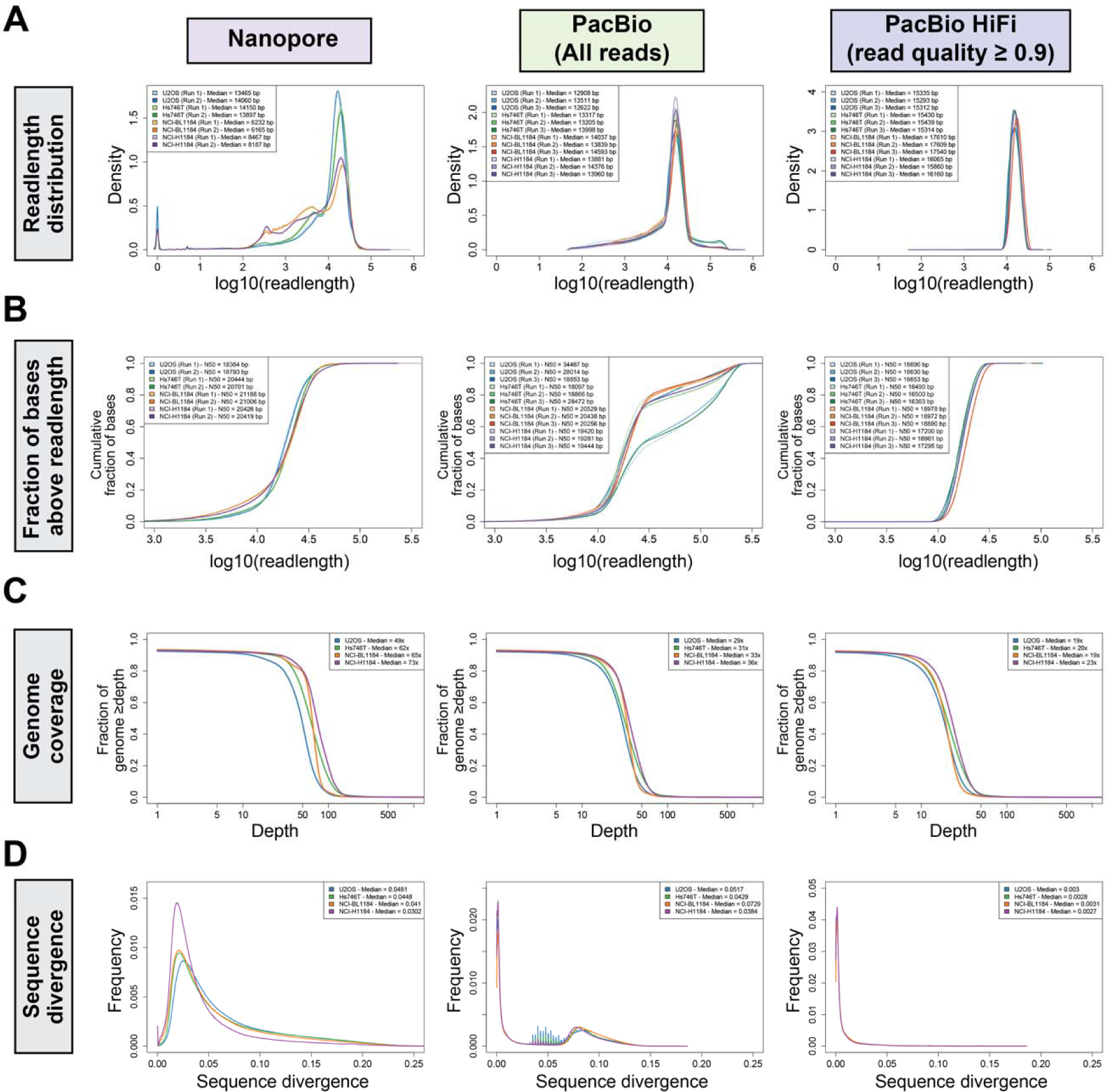
Sequencing quality of long-read genome sequencing datasets generated as part of this study. Sequencing quality metrics for long-read genome sequencing data generated using the Nanopore PromethION platform (2x flow cells per cell line), and the PacBio platforms (3x SMRT flow cells per cell line) are as indicated. The PacBio dataset was further divided into the full dataset consisting of all reads, and the PacBio HiFi dataset consisting of reads with a read quality ≥ 0.9. **(A)** Read length distribution of the Nanopore and PacBio long-read genome sequencing datasets generated in this study. Each sequencing run is indicated separately. The median read length for each run are also indicated in the legend for each plot. **(B)** The cumulative fraction of sequenced bases above a particular read length for each run is as indicated in the plots. The N50 (i.e. minimum read length at which half the bases were sequenced) for each run is also indicated in the legend of each plot. **(C)** Fraction of the genome sequenced above the stated sequencing depth for each sample is as indicated in the plots. The median sequencing coverage across the human genome is also indicated in the legend of each plot. **(D)** Distribution of sequence divergence of long-reads generated by each platform and for each platform is as indicated. Sequence divergence (i.e. how much each long-read differs from the reference genome) information for each long-read was extracted from long-reads aligned to the GRCh38 reference genome using minimap2.

**Figure S4.**
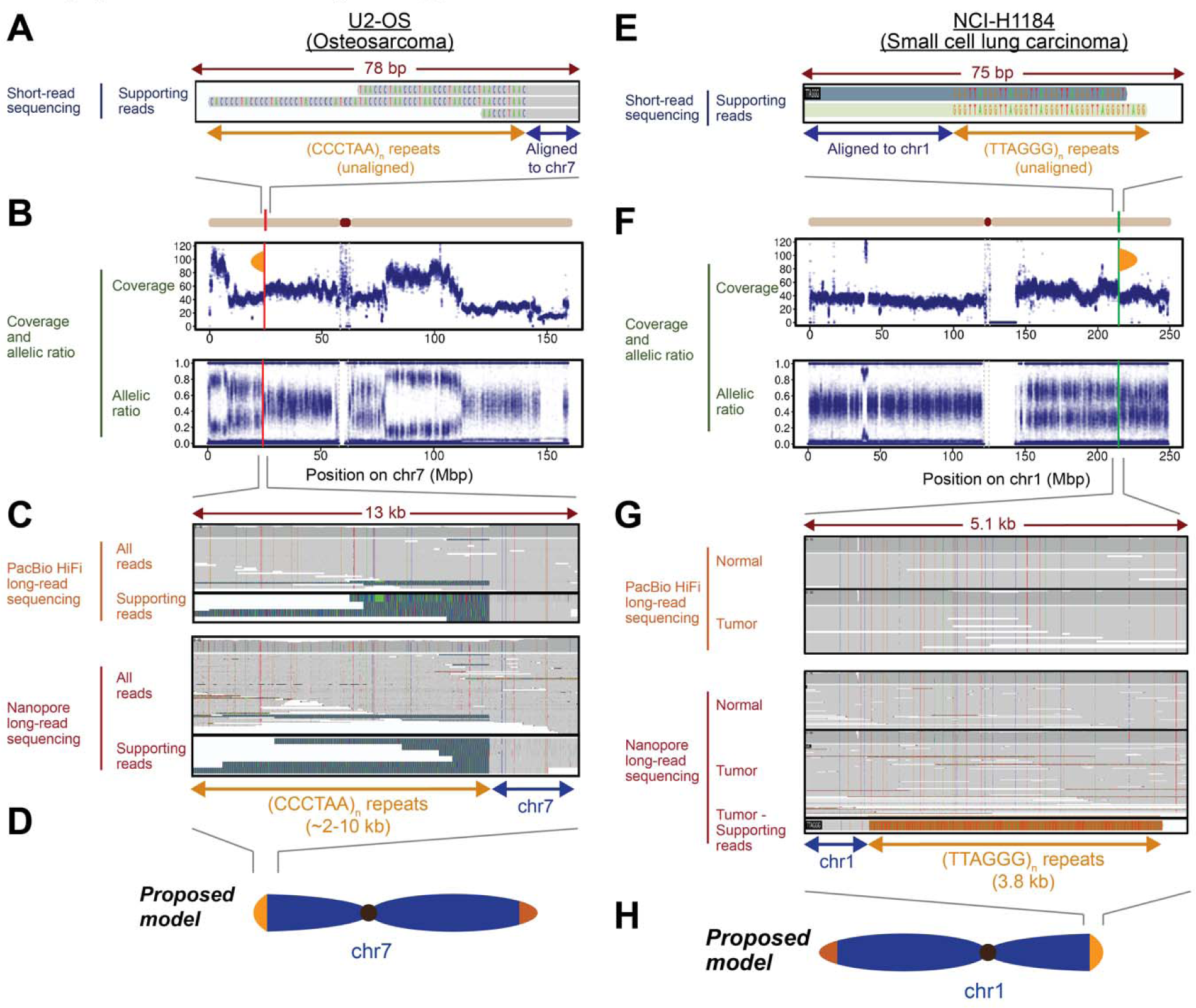
Additional examples of neotelomeres discovered using long-read genome sequencing, related to Figure 2. **(A-H)** Genomic analysis of telomere repeat alterations in the standard orientation that were detected **(A-D)** in the U2-OS osteosarcoma cell line at chr7:24,302,169, and **(E-H)** in the NCI-H1184 small cell lung carcinoma cell line at chr1:214,460,753, but not in the matched normal cell line (NCI- BL1184). **(A)** IGV screenshots of short-read genome sequencing data. Ectopic telomeric repeats (CCCTAA)_n_ are shown in gold. **(B)** Sequencing coverage and allelic ratios of chromosome 7. Orange semi-oval: site of the neotelomeric event. **(C)** IGV screenshots depicting long telomeric repeat sequences (TTAGGG)_n_ with PacBio HiFi (read quality ≥ 0.9) and Nanopore long-read sequencing at the site shown in (**A**). **(D)** Schematic of neotelomere location on chromosome 7p. **(E)** IGV screenshots of short-read genome sequencing data. Ectopic telomeric repeats (TTAGGG)_n_ are shown in gold. **(F)** Sequencing coverage and allelic ratios of chromosome 1. Orange semi-oval: site of the neotelomeric event. **(G)** IGV screenshots depicting long telomeric repeat sequences (CCCTAA)_n_ with PacBio HiFi (read quality ≥ 0.9) and Nanopore long-read sequencing at the site shown in (**E**). **(H)** Schematic of neotelomere location on chromosome 1q.

**Figure S5.**
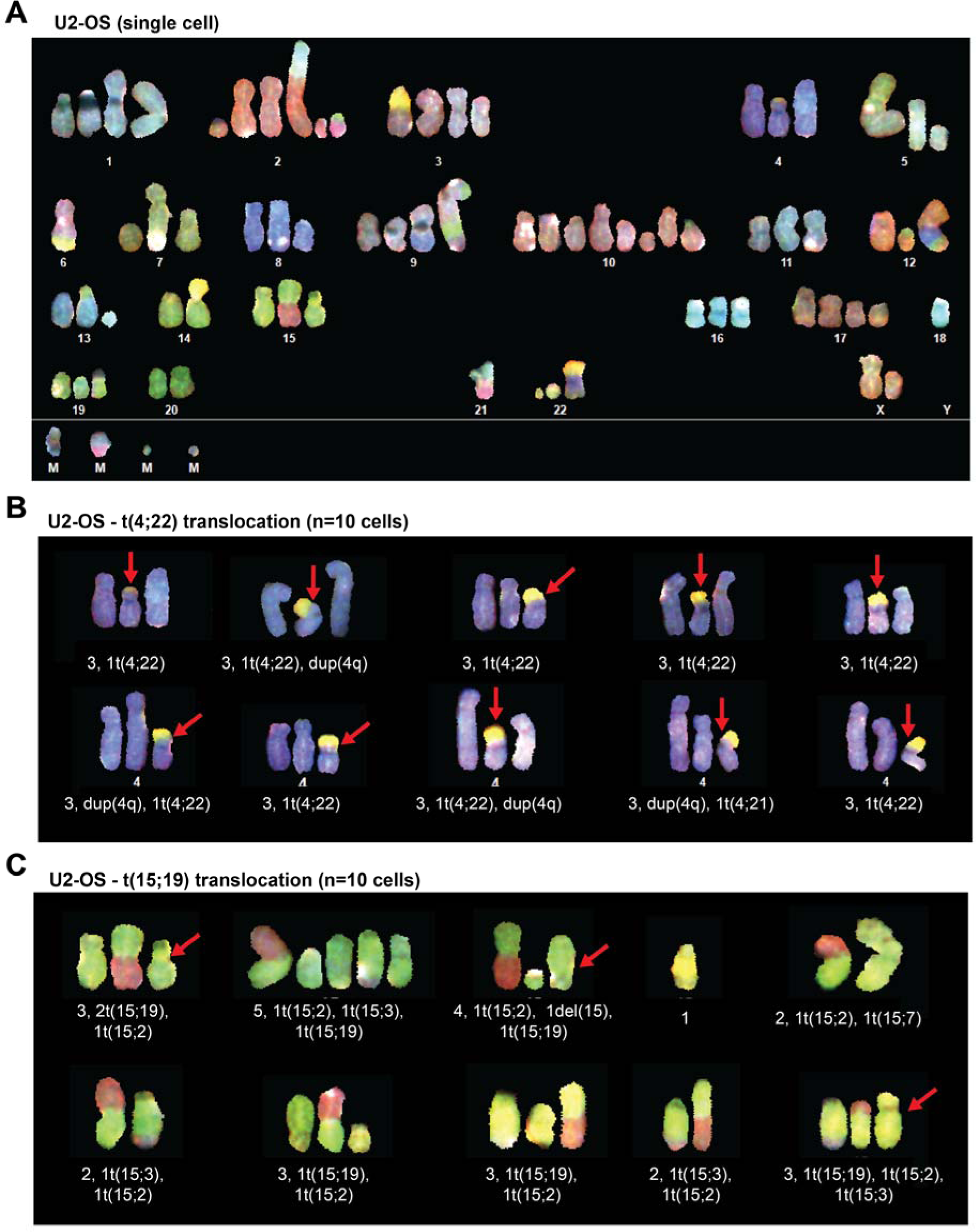
Degree of chromosomal heterogeneity between cells is chromosome specific. **(A)** Spectral karyogram of a representative U2-OS cell (Cell 01-01) analyzed in this study. Chromosomes observed were assigned to each of the 24 possible autosomes and sex chromosomes. Chromosomes that could not be assigned were labelled as marker chromosomes ‘M’. Spectral karyogram of **(B)** chromosome 4 with low levels of chromosomal heterogeneity and **(C)** chromosome 15 with high levels of chromosomal heterogeneity in ten cells assessed. Red arrows in **(B)** highlights the chromosome with translocation between chromosome 4 and 22. Red arrows in **(C)** highlights the chromosome with translocation between chromosome 15 and 19.

**Figure S6.**
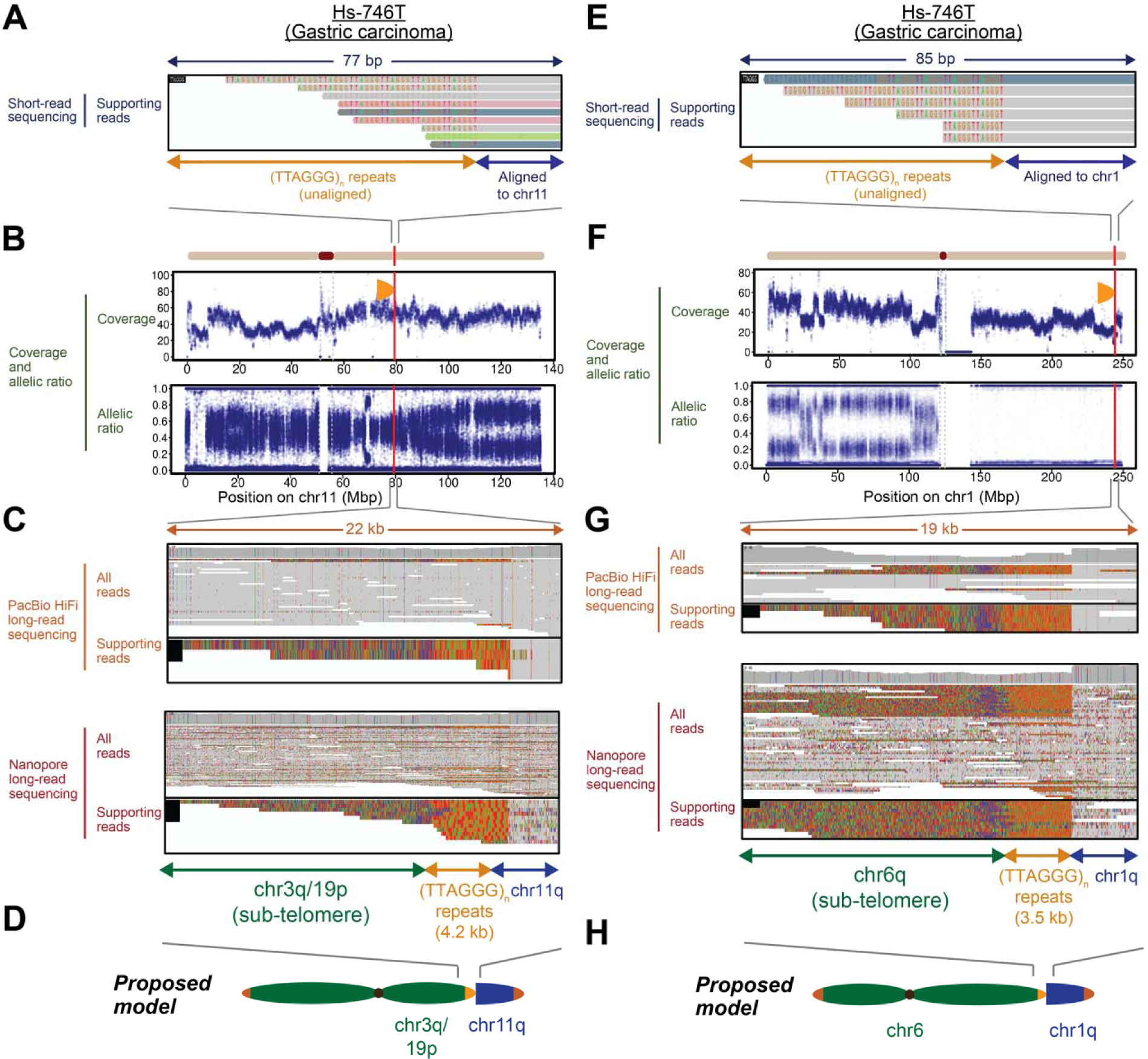
Additional examples of chromosomal arm fusion events revealed by long-read genome sequencing, related to Figure 3. **(A-H)** Genomic analysis of telomere repeat alterations in the inverted orientation that were detected in the Hs-746T gastric carcinoma cell line **(A-D)** at the site chr11:79,325,679, and **(E-H)** at the site chr1:244,201,717. **(A)** IGV screenshots of short-read genome sequencing data. Ectopic telomeric repeats (TTAGGG)_n_ are shown in color. **(B)** Sequencing coverage and allelic ratios of chromosome 11. Orange semi-oval: site of the ectopic telomere repeat sequence. **(C)** IGV screenshots of PacBio HiFi (read quality ≥ 0.9) and Nanopore long-read sequencing data at the site shown in (**A**). Ectopic telomeric repeats in the inverted orientation contained ∼4.2 kb of (TTAGGG)_n_ telomeric repeat sequences followed by chr3q/19p sub-telomeric sequences, indicative of a chromosomal arm fusion event of chr3q/19p to the site at chr11:79,325,679. **(D)** Schematic of telomere-spanning fusion event between chromosomes 3q/19p-ter and 11q. **(E)** IGV screenshots of short-read genome sequencing data. Ectopic telomeric repeats (TTAGGG)_n_ are shown in color. **(F)** Sequencing coverage and allelic ratios of chromosome 1. Orange semi-oval: site of the ectopic telomere repeat sequence. **(G)** IGV screenshots of PacBio HiFi (read quality ≥ 0.9) and Nanopore long-read sequencing at the site shown in (**E**). Ectopic telomeric repeats in the inverted orientation contained ∼3.5 kb of (TTAGGG)_n_ telomeric repeat sequences followed by chr6q sub-telomeric sequences, indicative of a chromosomal arm fusion event of chr6q to the site at chr1:244,201,717. **(H)** Schematic of telomere-spanning fusion event between chromosomes 6q-ter and 1q.

**Figure S7.**
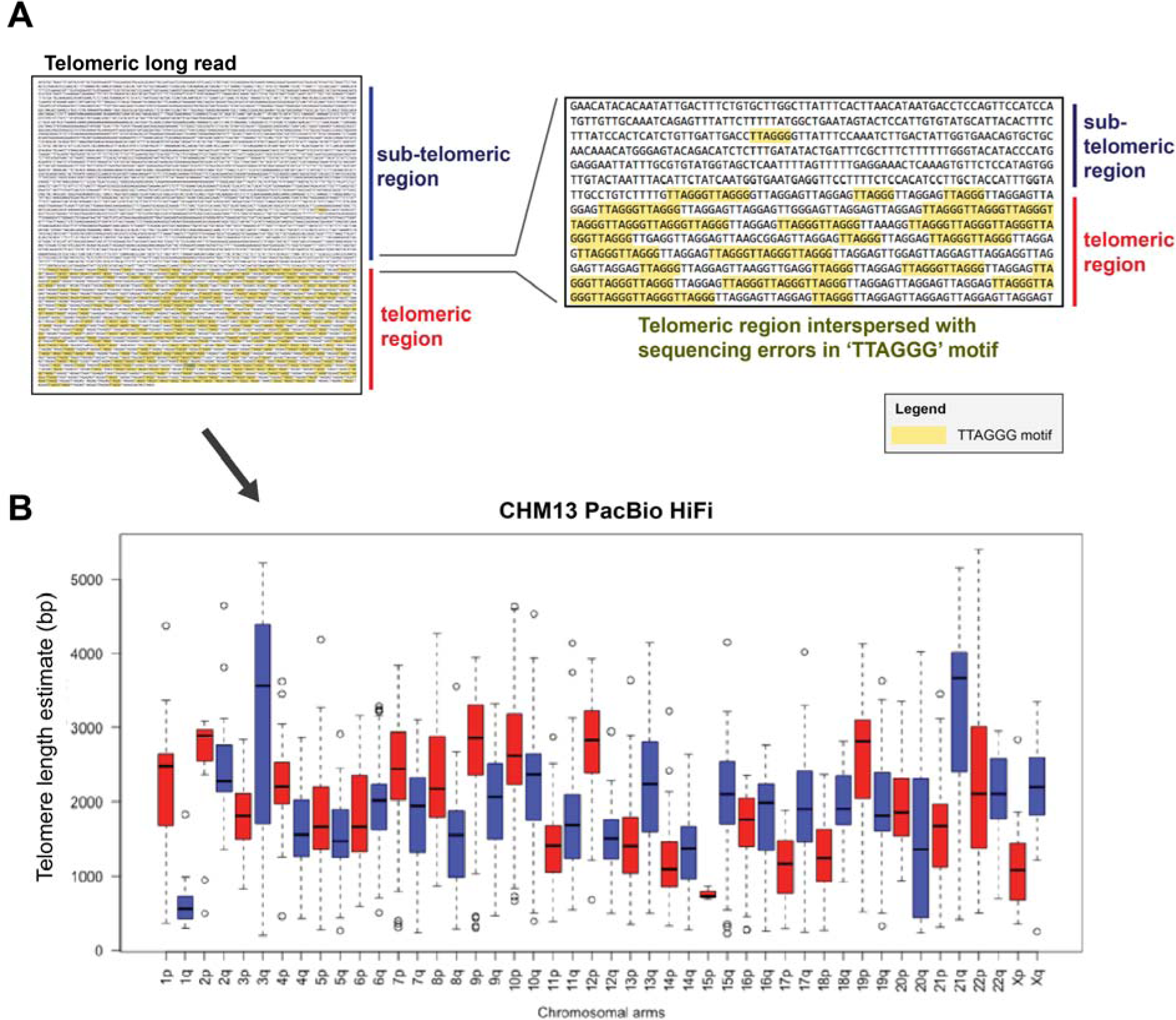
Estimation of telomere length from telomeric long-reads. **(A)** Schematic depicting how the telomeric region from a single telomeric long-read is defined. The TTAGGG motif on a single telomeric long-read is highlighted in yellow on the left, and a concentration of telomeric repeats can be observed towards the end of the telomeric long-read. The telomeric region from the single long-read can then be defined to estimate telomere length on the single long-read. A zoomed-in view of the boundary between the sub-telomeric and telomeric region is provided on the right. **(B)** Telomere length estimate for each chromosomal arm in the CHM13 cell line determined using PacBio HiFi long-read genome sequencing.

**Figure S8.**
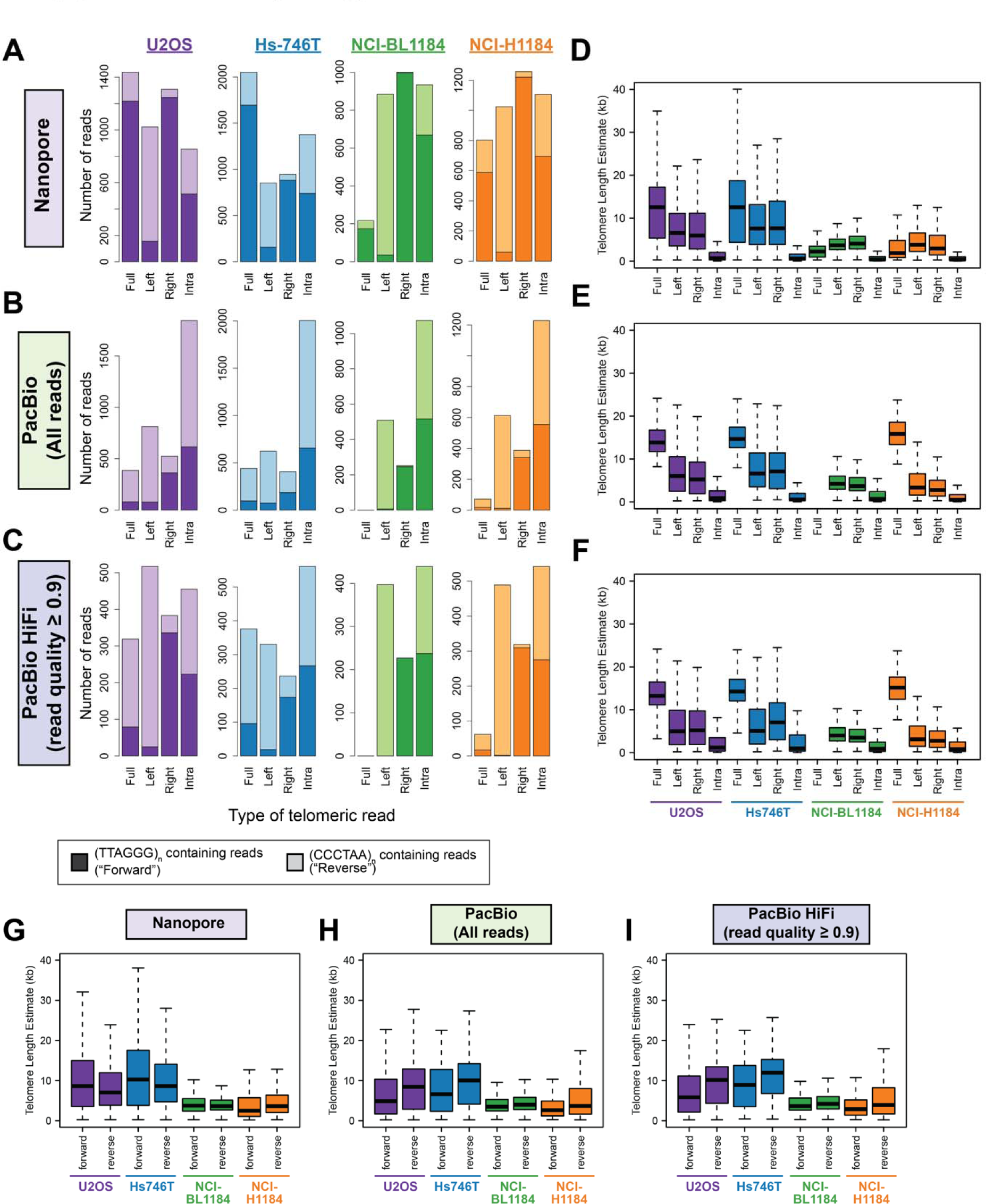
Telomere length estimates for the cell lines sequenced in this study using different sequencing platforms. **(A)** Number of telomeric reads of each class in the long-read datasets generated in this study. Long-reads containing telomeric repeats were split into four different classes depending on where telomeric repeats were observed in the long-read. These four classes are: Full – Long-reads that contains telomeric repeat sequences end-to-end, Left – Long-reads that contains telomeric repeat sequences on the left edge of the long-read, Right – Long-reads that contains telomeric repeat sequences on the right edge of the long-read, and Intra – Long-reads that contains telomeric repeat sequences in the middle of the single long-read. The type of telomeric repeat sequences observed is also further indicated (i.e. if the reads contain (TTAGGG)_n_ or (CCCTAA)_n_ repeats). Results for the four cell lines sequenced in this study by Nanopore, PacBio (All reads), or PacBio HiFi (read quality ≥ 0.9) sequencing are as indicated. **(D-F)** Telomere length estimates for the four classes of telomeric reads in the four cell lines sequenced. Results for each of the sequencing platforms: **(D)** Nanopore, **(E)** PacBio (All reads) and **(F)** PacBio HiFi (read quality ≥ 0.9) are as indicated. **(G-I)** Telomere length estimates for telomeric reads derived from either the “forward” strand (i.e. containing (TTAGGG)_n_ repeats) or “reverse” strand (i.e. containing (CCCTAA)_n_ repeats) are as indicated. Results for each of the sequencing platforms: **(G)** Nanopore, **(H)** PacBio (All reads) and **(I)** PacBio HiFi (read quality ≥ 0.9) are as indicated.

**Figure S9.**
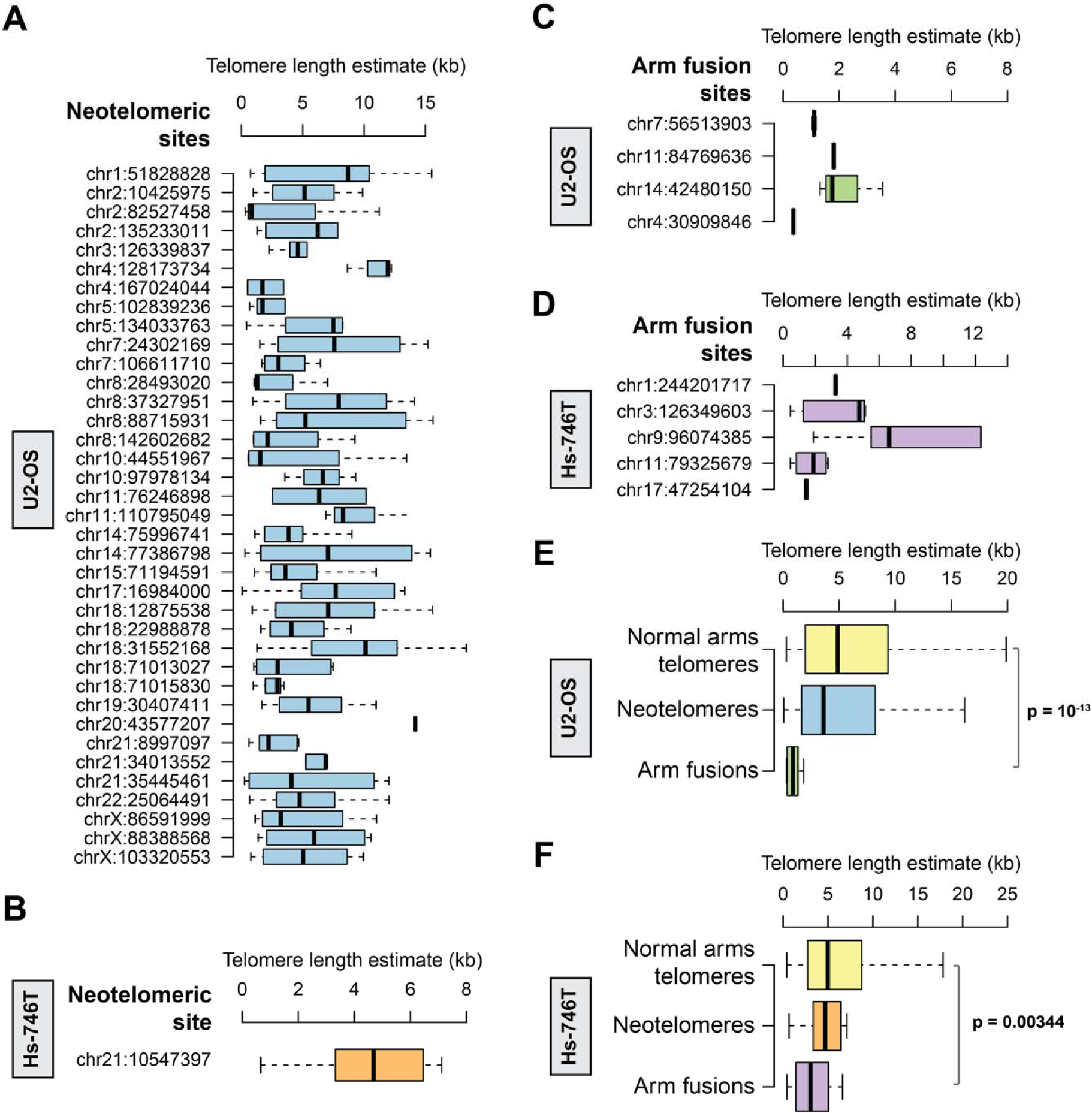
Length of telomeric repeats at neotelomeres and chromosomal arm fusion events as estimated using PacBio HiFi sequencing, related to Figure 4. The length of telomeric repeats on each long-read was estimated from these telomeric repeat signal profiles. Boxplots depicting the distribution of telomere length found at each neotelomere assessed by PacBio HiFi for the **(A)** U2-OS and (B) Hs-746T cell lines. Boxplot depicting length of telomeric repeats assessed using PacBio HiFi for each chromosomal arm fusion event in the **(C)** U2-OS and **(D)** Hs-746T cell lines. Note: telomere length for neotelomeres and normal chromosomal arms were only estimated using long-reads reads that start or end in telomeric repeats, while length of telomeric repeats at chromosomal arm fusions were estimated using long-reads with telomeric repeats in the middle of the read. Aggregated telomeric length of all long-reads at the normal chromosomal arms (p- and q-arms), neotelomeres, and chromosomal arm fusion events in the **(E)** U2-OS and **(F)** Hs-746T cell lines. P-values indicated in the plots were calculated using the two-sided Wilcoxon Rank Sum test.

**Figure S10.**
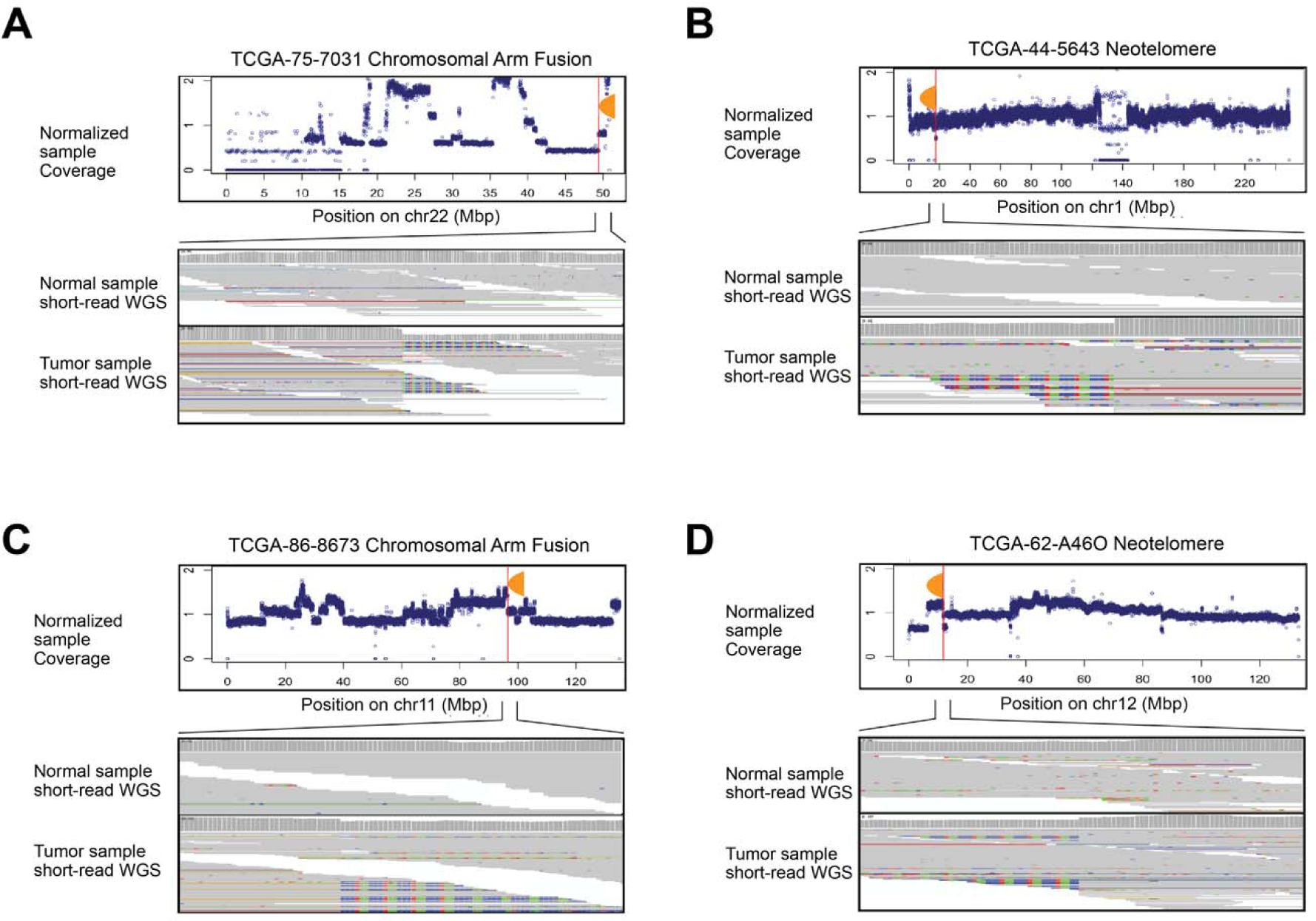
Representative examples of neotelomeres and chromosomal arm fusion events detected in patients with lung adenocarcinoma. **(A-D)** Normalized tumor sequencing coverage of chromosomes with neotelomeres and chromosomal arm fusion events predicted by TelFuse analysis of short-read genome sequencing are as depicted. Sequencing coverage of the tumor was normalized to the matched normal sample (Methods). IGV screenshots of the tumor and matched normal samples with neotelomeres and chromosomal arm fusion events are also as indicated. The sites and samples represented in the plot are **(A)** the putative chromosomal arm fusion site chr22:49,418,106 in TCGA-75-7031, **(B)** the putative neotelomere site chr1:17,644,075 in TCGA-44-5643, **(C)** the putative chromosomal arm fusion site chr11:96,570,712 in TCGA-86-8673, and **(D)** the putative neotelomere site chr12:11,696,012 in TCGA-62- A46O.

**Figure S11.**
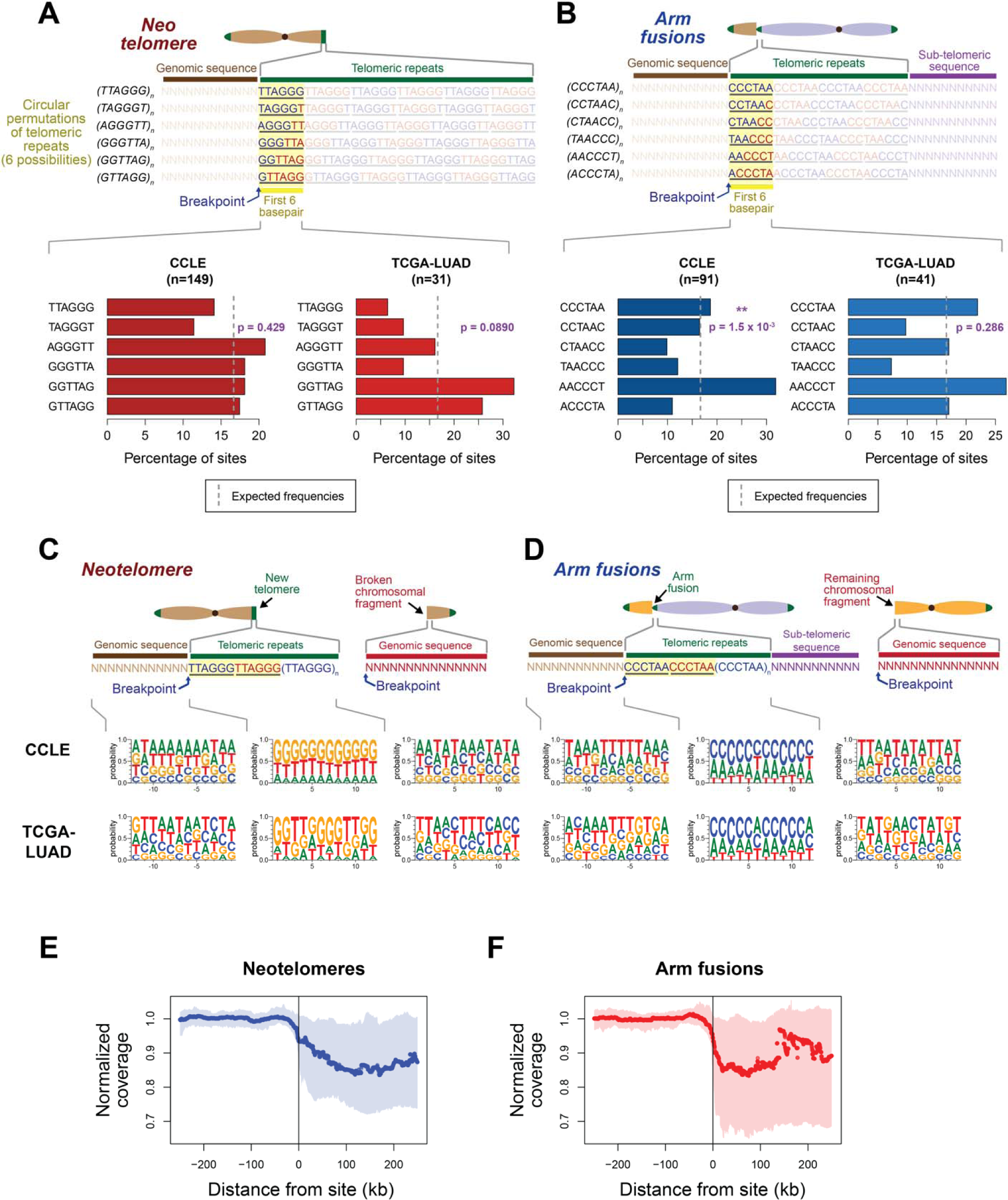
Little to no sequence preference associated with neotelomeres and chromosomal arm fusion events. **(A-B)** The first 6 base-pairs of each stretch of telomeric repeat sequence at each **(A)** neotelomere or **(B)** chromosomal arm fusion event was assessed and classified into one of six possible circular permutations representing the telomeric repeat sequence. **(A)** (top) Schematic illustrating the telomeric repeat sequences that are found directly after a breakpoint, and at a neotelomere. The first 6 base-pairs of the neotelomere after the breakpoint can occur in anyone of six possible circular permutations of the TTAGGG sequence. (bottom) Bar plots depicting the frequency of each six possible circular permutations observed on the first 6 base-pairs of the neotelomere in the CCLE and TCGA-LUAD cohorts. **(B)** (top) Schematic illustrating the telomeric repeat sequences that are found directly after a breakpoint, and at a chromosomal arm fusion site. The first 6 base-pairs of the chromosomal arm fusion after the breakpoint can occur in anyone of six possible circular permutations of the TTAGGG sequence. (bottom) Bar plots depict the frequency of each six possible circular permutations observed on the first 6 base-pairs of the chromosomal arm fusions observed in the CCLE and TCGA-LUAD cohorts. p-values in **(A)** and **(B)** were calculated using the chi-squared test under the assumption that all six circular permutations are expected to be observed at the same frequency. The expected frequencies are indicated by a grey dotted line, and the number of events assessed for each cohort is indicated in the header of each plot. **(C-D)** Sequence logo plot representing the frequencies of nucleotides observed near the breakpoints of neotelomere and chromosomal arm fusion events. **(C)** (top) Schematic of the neotelomere, and the three main regions (genomic region flanking the neotelomere, telomeric repeats corresponding to the neotelomeres, and genomic region of the broken chromosomal fragment that was detached from the neotelomere) associated with these events. (bottom) Logo plots representing frequencies of nucleotides in the three main regions around a neotelomeric event. **(D)** (top) Schematic of a chromosomal arm fusion event, and the four main regions around the breakpoint of the chromosomal arm fusion event (genomic region flanking the arm fusion event, telomeric repeats corresponding to the chromosomal arm that fused to this site, sub-telomeric region of the arm that underwent fusion, and the genomic region of the remaining chromosomal fragment that was detached following the chromosomal arm fusion event). (bottom) Frequency of nucleotides in the three regions around the breakpoint of a chromosomal arm fusion event. (E-F) Coverage profiles in the ± 200kb region surrounding a **(E)** neotelomere or **(F)** telomere fusion event in the CCLE cohort. The line depicts the median coverage observed across all sites, while the shaded area represents the interquartile range.

**Figure S12.**
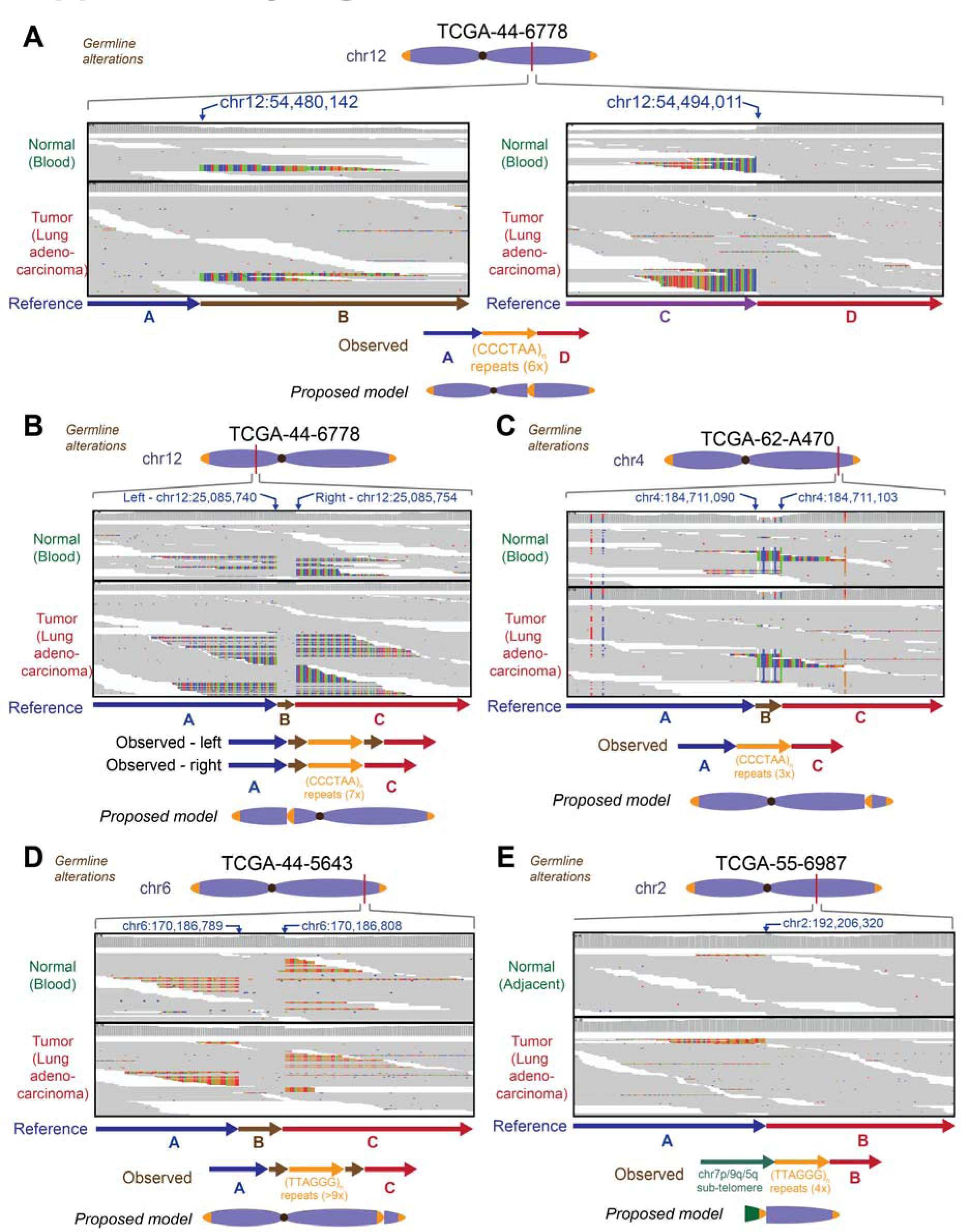
Putative germline ectopic telomeric events observed in lung adenocarcinoma tumor samples from patients. **(A)** Ectopic telomeric repeat sequences in the inverted orientation at the site chr12:54,480,142, and in the standard orientation at the site chr12:54,494,011 in both the normal (blood) and tumor (lung adenocarcinoma) sample for the patient TCGA-44-6778. IGV screenshots depicting these observations are as indicated. These observations point to a model where ∼6x (CCCTAA)_n_ repeats have integrated into this locus at chr12q, coupled with a deletion of regions B and C indicated in the figure. **(B)** Ectopic telomeric repeat sequences in the standard orientation were found at the site chr12:25,085,740, and in the inverted orientation at the site chr12:25,085,754 in both the normal (blood) and tumor (lung adenocarcinoma) sample for the patient TCGA-44-6778. IGV screenshots depicting these observations are as indicated. These observations point to a model where ∼7x (CCCTAA)_n_ repeats have integrated into this locus at chr12p, coupled with a duplication of region B for the event on the left. The event on the right represents the insertion of the telomeric repeats without duplication of region B. **(C)** Ectopic telomeric repeat sequences in the inverted orientation at the site chr4:184,711,090, and in the standard orientation at the site chr4:184,711,103 in both the normal (blood) and tumor (lung adenocarcinoma) sample for the patient TCGA-62-A470. IGV screenshots depicting these observations are as indicated. These observations point to a model where ∼3x (CCCTAA)_n_ repeats have integrated into this locus at chr4q, coupled with a deletion of region B found on the reference genome. **(D)** Ectopic telomeric repeat sequences in the inverted orientation was found at the site chr6:170,186,789, and in the standard orientation at the site chr6:170,186,808 in both the normal (blood) and tumor (lung adenocarcinoma) sample for the patient TCGA-44-5643. IGV screenshots depicting these observations are as indicated. These observations point to a model where >9x (TTAGGG)_n_ repeats have integrated into this locus at chr6q, coupled with a duplication of region B in the reference genome. **(E)** Ectopic telomeric repeat sequences in the inverted orientation was found at the site chr2:192,206,320 in both the normal (adjacent lung tissue) and tumor (lung adenocarcinoma) sample for the patient TCGA-55-6987. IGV screenshots depicting these observations are as indicated. These observations point to a model where 4x (TTAGGG)_n_ repeats have integrated into this locus at chr2q, together with sub-telomeric sequences corresponding to either chr7q/9q/5q, suggesting that a chromosomal arm fusion event has potentially occurred here.

**Figure S13.**
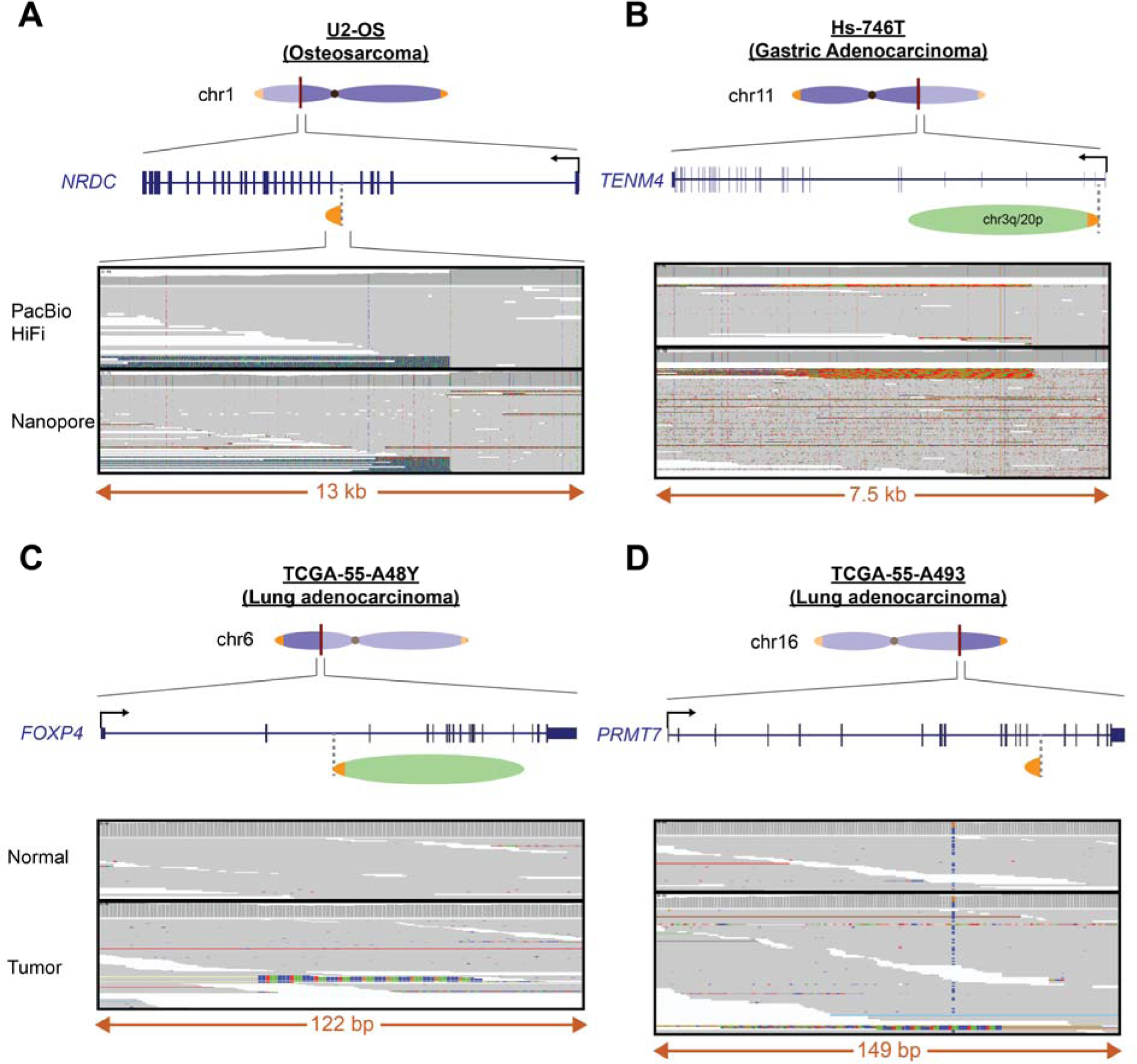
Additional examples of neotelomeric and chromosomal arm fusion events which led to gene disruptions, related to Figure 6. **(A-B)** Schematic and IGV screenshots depicting gene disrupting events caused by neotelomeres or chromosomal arm fusion events in cancer cell lines. These were observed in (A) the U2-OS osteosarcoma cell line where a neotelomere could be observed in the middle of the *NRDC* gene at the site chr1:51,828,828, and **(B)** the Hs-746T gastric adenocarcinoma cell line where a chromosomal arm fusion event could be observed within the *TENM4* gene at the site chr11:79,325,679. **(C-D)** Somatic neotelomere and chromosomal arm fusion events observed in primary lung adenocarcinoma tumor samples. These were observed in the tumor sample of **(C)** patient TCGA-55-A48Y in the middle of the *FOXP4* gene at the position chr6:41,573,027 where a putative chromosomal arm fusion is observed, and in **(D)** patient TCGA-55-A493 in the *PRMT7* gene at the position chr16:68,349,160, where a putative neotelomere could be observed.

### Supplementary Table Legends

**Table S1 SRA accession numbers of short-read genome sequencing data for cancer cell lines analyzed in this study.**

**Table S2 Detailed information of ectopic telomeric sites identified in cancer cell lines analyzed in this study.** Sites indicated in this table has perfect telomeric repeat sequences on the first 12 base-pairs of the event.

**Table S3 Detailed information of ectopic telomeric sites identified in cancer cell lines analyzed in this study without perfect telomeric repeat sequences on the first 12 base-pairs.** Sites indicated in this table do not have perfect telomeric repeat sequences on the first 12 base-pairs of the event but contains at least 12 base-pairs of telomeric repeat sequences within the soft-clipped sequences.

**Table S4 Sequencing statistics of each long-read genome sequencing run generated for this study.**

**Table S5 Sequencing statistics for each sample analyzed by long-read genome sequencing for this study.** Multiple runs for the same sample were aggregated into a single dataset, and their corresponding sequencing metrics are as indicated.

**Table S6 Sites assessed, and validation status as determined by long-read genome sequencing.**

**Table S7 Spectral karyotyping results of ten U2-OS cells.**

**Table S8 Genomic Data Commons accession numbers for TCGA Lung adenocarcinoma patient samples analyzed in this study.**

**Table S9 Detailed information of ectopic telomeric sites identified in tumor samples in the cohort of lung adenocarcinoma samples analyzed.**

**Table S10 Detailed information of ectopic telomeric sites identified in normal samples in the cohort of lung adenocarcinoma samples analyzed.**

